# Quantitative proteomics combined with network pharmacology analysis unveils the biological basis of Schisandrin B in treating diabetic nephropathy

**DOI:** 10.1101/2023.01.19.524316

**Authors:** Jianying Song, Bo Zhang, Xudong Lyu, Huiping Zhang, Wenbo Cheng, Peiyuan Liu, Jun Kang

## Abstract

**Background:** Diabetic nephropathy (DN) is a major complication of diabetes. Schisandrin B (Sch) is a natural pharmaceutical monomer that was shown to prevent kidney damage caused by diabetes and restore its function. However, there is still a lack of comprehensive and systematic understanding of the mechanism of Sch treatment in DN.

**Objective:** We aim to provide a systematic overview of the mechanisms of Sch in multiple pathways to treat DN in rats.

**Methods:** Streptozocin was used to build a DN rat model, which was further treated with Sch. The possible mechanism of Sch protective effects against DN was predicted using network pharmacology and was verified by quantitative proteomics analysis.

**Results:** High dose Sch treatment significantly downregulated fasting blood glucose, creatinine, blood urea nitrogen, and urinary protein levels and reduced collagen deposition in the glomeruli and tubule-interstitium of DN rats. The activities of superoxide dismutase (SOD) and plasma glutathione peroxidase (GSH-Px) in the kidney of DN rats significantly increased with Sch treatment. In addition, the levels of IL-6, IL-1β, and TNF-α were significantly reduced in DN rats treated with Sch. 11 proteins that target both Sch and DN were enriched in pathways such as MAPK signaling, PI3K-Akt signaling, renal cell carcinoma, gap junction, endocrine resistance, and TNF signaling. Furthermore, quantitative proteomics showed that Xaf1 was downregulated in the model vs. control group and upregulated in the Sch-treated vs. model group. Five proteins, Crb3, Tspan4, Wdr45, Zfp512, and Tmigd1, were found to be upregulated in the model vs. control group and downregulated in the Sch vs. model group. Three intersected proteins between the network pharmacology prediction and proteomics results, Crb3, Xaf1, and Tspan4, were identified.

**Conclusion:** Sch functions by relieving oxidative stress and the inflammatory response by regulating Crb3, Xaf1, and Tspan4 protein expression levels to treat DN disease.

## 1. INTRODUCTION

Diabetic nephropathy (DN) is one of the major complications of diabetes and a major cause of end-stage renal failure, with about 30% of diabetic patients deteriorating into DN [1]. Early symptoms include increased glomerular filtration capacity and the appearance of microalbumin in urine. If not treated, glomerular atrophy, basement membrane thickening, and extracellular matrix accumulation may lead to chronic renal insufficiency [2]. Deteriorating even further into clinical proteinuria, the renal function will experience a sharp decline that gives rise to end-stage renal failure [3], seriously affecting the quality of life of patients. At present, the treatment methods for DN mainly encompass controlling blood glucose levels and regulating lipid metabolism with antihypertensive drugs [4, 5] and vitamins [6]. However, existing treatment methods have failed to completely prevent the deterioration of DN [7]. Therefore, new drugs or treatments that could improve the renal function of DN patients and alleviate the progression of the disease have attracted much attention.

The immune and inflammatory mechanisms play a pivotal role in the development of DN, which is considered to be a chronic inflammatory disease [8]. Several types of cells, such as monocytes, macrophages, and lymphocytes, as well as chemokines and cytokines, are involved in this process [9, 10]. Among them, it is well known that interleukin 6 (IL-6), interleukin 1 beta (IL-1β), and tumor necrosis factor-alpha (TNF-α) are related to the progression of DN, due to their ability to induce kidney cells to produce various chemokines, triggering the next round of inflammatory responses [11, 12]. Diabetic patients often suffer from obesity, insulin resistance, and increased oxidative stress. It was also found in diabetic patients that the levels of heat shock protein 60 and free fatty acids were increased, the expression of TLR4 was significantly increased, and the downstream molecule NF-κB translocated into the nucleus. This activated the TLR4/NF-κB inflammatory signal transduction pathway, which promoted the transcription of downstream inflammation-related genes, inducing the production of a large number of cytokines and inflammatory mediators, which further led to renal inflammatory damage [13, 14].

Schisandrin B (Sch) is a natural active lignin compound extracted and isolated from the traditional Chinese medicine *Schisandra chinensis*. Sch was shown to exhibit significant antioxidant, anti-inflammatory, and anti-apoptotic effects and to support the health of organs such as the heart [15-17], brain [18, 19], liver [20], and kidney [21]. Sch was shown to be involved in the renal protection of type 2 DN rats by regulating the HMGB1/TLR4 signaling pathway [22]. Sch improved renal function by intervening in the Nrf2/ARE signaling pathway, thus exerting a protective effect on rats with nephrotic syndrome [23]. Sch also alleviated hyperglycemia-induced renal injury by suppressing excessive inflammation and oxidative stress through maintenance of urine creatinine and albumin levels in a diabetic mouse model [24]. Despite the exceptional curative potential and effects on DN, the overall systematic understanding of the molecular mechanism of Sch in the treatment of DN disease is still lacking.

As the globalization movement and digitalization become the mainstream of modern research, the combination of bioinformatics with traditional experimental methods has received much attention and enthusiasm. Network pharmacology is a new discipline based on the theory of system biology, which analyzes a network of biological systems and selects specific signal nodes for molecular design of multiple drug targets [25, 26]. Traditional Chinese medicine (TCM) network pharmacology, which is firstly proposed by Li [27, 28], provide a new systematic way to understanding TCM and is widely used in exploring mechanism underlying natural products [29, 30]. It is a useful and comprehensive tool to elucidate complicated relationships among drugs, gene targets, and diseases based on building a network. Network pharmacology emphasizes the regulation of signaling pathways in multiple ways to improve the therapeutic effect of drugs and reduce side effects, so as to improve the success rate of clinical trials of new drugs and reduce the cost of drug research [31-35].

In this article, high-fat feed combined with streptozocin (STZ) injection was used to establish a DN rat model. The effect of Sch treatment on differential expression of proteins in the kidneys of the DN rats was explored and clarified by network pharmacology and proteomics for a systematic and comprehensive understanding of the molecular mechanism of Sch treatment of DN disease (Scheme 1).

## 2. MATERIALS AND METHODS

### 2.1. Reagents

Sch was purchased from Shanghai Yuanye Bio-Technology Co., Ltd. (61281-37-6, Shanghai, China). Irbesartan was purchased from Lunan Beite Pharmaceutical Co., Ltd. (Batch No. 36160901, specification: 40 mg/tablet, Shandong, China).

### 2.2. Modeling

Fifty healthy specific-pathogen free male Sprague-Dawley rats, 6–8 weeks old, weighing 200±20 g, were provided by Beijing HFK Bioscience Co., Ltd. (quality certificate No. 110322210100331916). The feeding environment was maintained at 25±2°C, the relative humidity was 50±15%, there was a 12-h light and dark cycle, and the rats had free access to food and water. The experimental conditions were improved and optimized based on Azushima’s research [36]. The experiment was approved by the Ethics Committee of Tianjin University.

After 1 week of adaptive feeding, 10 rats were randomly chosen as the negative control group, and the other 40 rats were fed a high sugar and high fat diet (10% lard, 20% sucrose, 2.5% cholesterol, and 67.5% conventional diet). Intraperitoneal injection of STZ at 30 mg/kg [1% STZ solution prepared with 0.1 mmol/L citric acid-sodium citrate buffer (pH 4.4)] was performed at the end of the 7th week. After 72 h, blood samples from the tail vein were randomly collected to test blood glucose content. The modeling of diabetic rats was regarded as successful if the blood glucose concentration exceeded 16.7 mmol/L. Diabetic rats were fed for another week, and 24 h urine was collected using metabolic cages after 1 week to measure urine protein content. The DN animal model was established with the standard of a blood glucose concentration exceeding 16.7 mmol/L, positive urine glucose, and positive urine protein. During modeling, the weight of the rats in each group was recorded every 2 weeks.

Successfully modeled rats were randomly divided into a model group, a positive control group, an Sch low dose group, and an Sch high dose group, with 10 rats in each group. The positive control group was administered irbesartan at 30 mg/kg [37], the Sch low dose group at 20 mg/kg, and the Sch high dose group at 40 mg/kg. The negative control group and model group were administered the same volume of distilled water by intragastric administration for 4 continuous weeks. During the experiment, the rats were fed a standard diet and had free access to drinking water. Food was withheld for 12 h every Saturday night to collect blood from the tail vein to measure the fasting blood glucose at 8:00 a.m. on Sunday.

### 2.3. Biochemical index analysis

Four weeks after the Sch intervention, 24 h urine collections for each rat from each group were performed by using metabolic cages. The urine was centrifuged at 4000 g for 10 min, and the 24 h urinary albumin concentration was measured. The rats were anesthetized by an intraperitoneal injection of 50 mg/kg pentobarbital sodium, and the inner canthus blood was collected. The collected blood was centrifuged at 3000 r/min for 15 min, and serum was collected. The serum levels of serum creatinine and blood urea nitrogen (BUN) were determined by an automatic biochemical analyzer.

After blood collection, the rats were killed using cervical dislocation. Kidneys were removed immediately, and part of the rat kidney tissue was extracted, washed with pre-cooled phosphate-buffered saline three times, homogenized at a low temperature after cutting, centrifuged at 16,000 g for 10 min, and the supernatant was collected. The activities of superoxide dismutase (SOD) and plasma glutathione peroxidase (GSH-Px) and malondialdehyde (MDA) levels related to oxidative stress were measured according to the steps described in the kit.

### 2.4. ELISA

The levels of IL-6, IL-1β, and TNF-α in the renal tissue homogenate were determined by ELISA. First, 100 μL of IL-6, IL-1β, and TNF-α captured antibodies was added to ELISA plates and incubated overnight at room temperature. The plates were washed and blocked for 1 h, then washed, and 100 μL of rat kidney tissue homogenate was added. Three replicates were used for each sample. The plates were incubated at room temperature for 3 h and washed afterwards. The plates were incubated with the corresponding enzyme-conjugate secondary antibody for 30 min and washed afterwards. After adding the chromogenic solution, the absorbance was read at a wavelength of 405 nm, and the concentrations of IL-6, IL-1β, and TNF-α were calculated.

### 2.5. Histopathological staining of the kidney

The renal tissues of rats in each group were fixed with a 10% formalin solution for 24 h, washed with water for 20 min, dehydrated with an alcohol gradient, made transparent with xylene, embedded in paraffin, and sectioned into 5-μm slices for hematoxylin and eosin (HE) staining, Masson staining, and Sirius red staining in order to observe the basic pathological changes of the renal tissues in each group under a microscope.

### 2.6. Drug and disease target prediction

The Sch target proteins were predicted using the ChEMBL (www.ebi.ac.uk) and CTD databases (ctdbase.org/). The screening standard was as follows: confidence was 90% for ChEMBL, and interaction was “increases, decreases, or affects” for the CTD database. The DN disease targets were predicted using the GeneCards (www.genecards.org) and TCMSP (old.tcmsp-e.com) databases.

### 2.7. Quantitative proteomics

#### 2.7.1. Protein extraction and digestion

Highly abundant protein was first removed from the samples, and then, SDT buffer (4% SDS, 100 mM Tris-HCl, 1 mM DTT, pH 7.6) was used for sample lysis and protein extraction. The amount of protein was quantified using a BCA Protein Assay kit (Bio-Rad, USA). An appropriate amount of protein was removed from each sample and digested by trypsin according to the filter-aided sample preparation procedure [38]. The digested peptides of each sample were desalted on C18 cartridges [Empore™ SPE Cartridges C18 (standard density), bed I.D. 7 mm, volume 3 mL, Sigma], concentrated by vacuum centrifugation, reconstituted in 40 µL of 0.1% (v/v) formic acid, and measured at OD_280_.

#### 2.7.2. Tandem mass tag labeling

Each sample containing a 100 μg peptide mixture was labeled using tandem mass tag reagent according to the manufacturer’s instructions (Thermo Scientific). The negative control group samples were labeled C1, C2, and C3, the model group samples were labeled M1, M2, and M3, and the Sch-treated model group samples were labeled S1, S2, and S3, so that each group had three biological replicates, for nine samples in total.

#### 2.7.3. Grading

For RP grading, the labeled peptides of each group were mixed in equal amounts and fractionated by a high pH reversed-phase peptide fractionation kit (Thermo Scientific). The column was first balanced by acetonitrile and 0.1% trifluoroacetic acid (TFA), then the mixed labeled peptide samples were loaded and pure water was added, followed by low-speed centrifugation for desalting. Finally, the column-bound peptides were gradient eluted by a high pH gradient of an acetonitrile solution with successively increasing concentrations. Each eluted peptide sample was vacuum-dried and lyophilized with 12 μL 0.1% FA, and the concentration of the peptides was determined by the OD_280_.

For SCX grading, the labeled peptides of each group were mixed and graded using an AKTA Purifier 100. Buffer A was 10 mM KH_2_PO_4_, and 25% ACN, pH 3.0, and buffer B was 10 mM KH_2_PO_4_, 500 mM KCl, and 25% ACN, pH 3.0. The chromatographic column was balanced with buffer A, and samples were added into the chromatographic column for separation at a flow rate of 1 mL/min. The gradient of the liquid phase was as follows: the gradient of buffer B was 0% for 25 min, and then, the linear gradient of buffer B increased from 0% to 10% from 25 to 32 min; then, it increased 10%–20% from 32 to 42 min; then, it increased 20%–45% from 42 to 47 min; then, it increased 45%–100% from 47 to 52 min and then remained at 100% from 52 to 60 min. After 60 min, buffer B was reset to 0%. The absorbance value at 214 nm was monitored during elution. Eluted components were collected at 1 min intervals and lyophilized with C18 cartridges for desalting.

#### 2.7.4. LC-MS/MS analysis

Each sample was separated by an HPLC liquid system, Easy nLC, at a nanoliter flow rate. Buffer A was a 0.1% formic acid aqueous solution, and buffer B was a 0.1% formic acid acetonitrile aqueous solution (the acetonitrile solution was 84%). The chromatographic column was balanced with 95% buffer A, and the samples were loaded onto the loading column by an automatic sampler (Thermo Scientific Acclaim PepMap100, 100 μm × 2 cm, nanoViper C18). The separation was performed on a Thermo Scientific EASY Column (10 cm, ID 75 μm, particle size 3 μm, C18-A2) at a flow rate of 300 nL/min. The samples were separated by chromatography and analyzed by a Q-EXactive mass spectrometer. The detection method was positive ion, the scanning range of the parent ion was 300–1800 m/z, the resolution of the first-level mass spectrometry was 70,000 at 200 m/z, the automatic gain control target was 1E6, the maximum IT was 50 ms, and the dynamic exclusion time was 60.0 s. The mass charge ratios of peptides and peptide fragments were collected as follows: twenty fragment profiles (MS2 scan) were collected after each scan. The MS2 activation type was HCD, and the isolation window was 2 m/z. The secondary mass spectrometry resolution was 17,500 at 200 m/z. The normalized collision energy was 30 eV, and the underfill was 0.1%.

#### 2.7.5. Identification and quantitation of proteins

The MS raw data for each sample were searched using the MASCOT engine (Matrix Science, London, UK; version 2.2) embedded in Proteome Discoverer 1.4 software for identification and quantitative analysis.

### 2.8. Statistical method

The statistical software SPSS 20.0 was used to analyze the experimental data. Data were expressed as mean ± standard deviation (x ± s), a *t* test was applied for comparisons between two groups, and one-way analysis of variance (ANOVA) was applied for comparisons between multiple groups. *P* < 0.05 was considered as a statistically significant difference.

## 3. RESULTS

### 3.1. Sch effects on blood glucose, renal function, and pathological manifestations in DN rats

After treating DN rats with Sch, the body weight of the model group decreased significantly (*P* < 0.01) compared to the negative control group, and the body weight of the positive control group, Sch low dose group, and Sch high dose group increased significantly (*P* < 0.01, *P* < 0.05, and *P* < 0.001, respectively) compared to the model group (Figure 1A).

**Figure 1.**
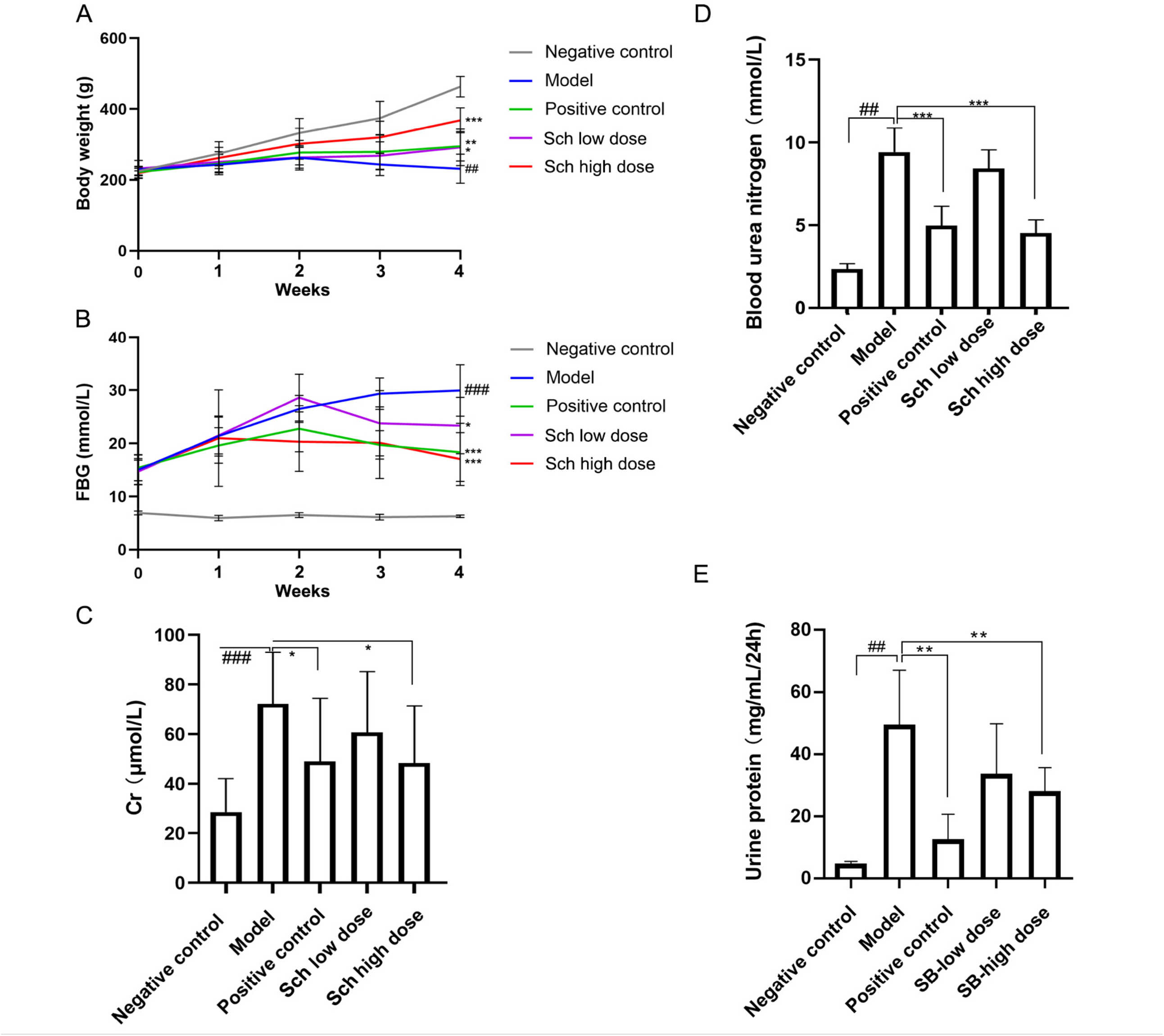
Sch treatment alleviated hyperglycemia and lowered creatinine, blood urea nitrogen, and urine protein in diabetic nephropathy (DN) rats. A. Sch treatment decreased the body weight of DN rats. B. Sch treatment decreased the fasting blood glucose in DN rats. C. Concentration of creatinine was decreased in Sch-treated DN rats. D. Blood urea nitrogen was decreased in Sch-treated DN rats. E. Urine protein was decreased in DN rats after Sch treatment. Negative control, model, positive control, Sch low dose, Sch high dose groups (n = 10 per group). Data are presented as the mean ± SD. ##: *P* < 0.01 as compared to the control group; ^*^: *P* < 0.05 as compared to the model group; ^**^: *P* < 0.01 as compared to the model group. ^***^: *P* < 0.001 as compared to the model group.

The concentrations of fasting blood glucose were above 16.7 mmol/L in all rats 72 h after STZ injection. After 4 weeks of Sch feeding, the fasting blood glucose in the Sch low dose group and Sch high dose group decreased significantly (*P* < 0.05 and *P* < 0.001, respectively, Figure 1B) compared to the model group. Renal function tests revealed that the creatinine, BUN, and 24 h urinary protein levels in the model group all increased significantly (*P* < 0.001, *P* < 0.01, and *P* < 0.001, respectively) compared to the negative control group. Irbesartan and high dose Sch treatments significantly reduced the levels of creatinine (*P* < 0.05 and *P* < 0.05, respectively) (Figure 1C), BUN (*P* < 0.001 and *P* < 0.001, respectively) (Figure 1D), and 24 h urinary protein (*P* < 0.001 and *P* < 0.001, respectively) (Figure 1E) in DN rats, while low-dose Sch treatment had no significant effect.

HE staining demonstrated that the glomerulus and renal tubules in the negative control group were normal, and no proliferation of mesangial cells or the mesangial matrix or inflammatory cell infiltration was observed. The model group showed focal degeneration and atrophy of renal tubules, slight thickening of the glomerular basement membrane, hyperplasia of mesangial cells, and steatosis of glomerular and renal tubules. Compared with the model group, the lesions in the experimental group were reduced, and the improvement was more obvious in the positive control group and Sch high dose group (Figure 2A). After Masson staining, collagen appeared blue under a light microscope. The glomerulus and renal tubules of rats in the negative control group were normal, and collagen expression did not increase, while collagen hyperplasia was observed in the mesangial area and basement membrane, and renal tubules were dilated in the model group. After Sch intervention, collagen deposition in the glomerulus and intertubular tissues of DN rats was significantly reduced (Figure 2B). Sirius red staining was used to indicate collagen deposition in the renal interstitium and vascular wall. The study showed that the glomerular arteriole wall of the model group was thickened, but the thickening degree of the high dose Sch treated group was weakened (Figure 2C).

**Figure 2.**
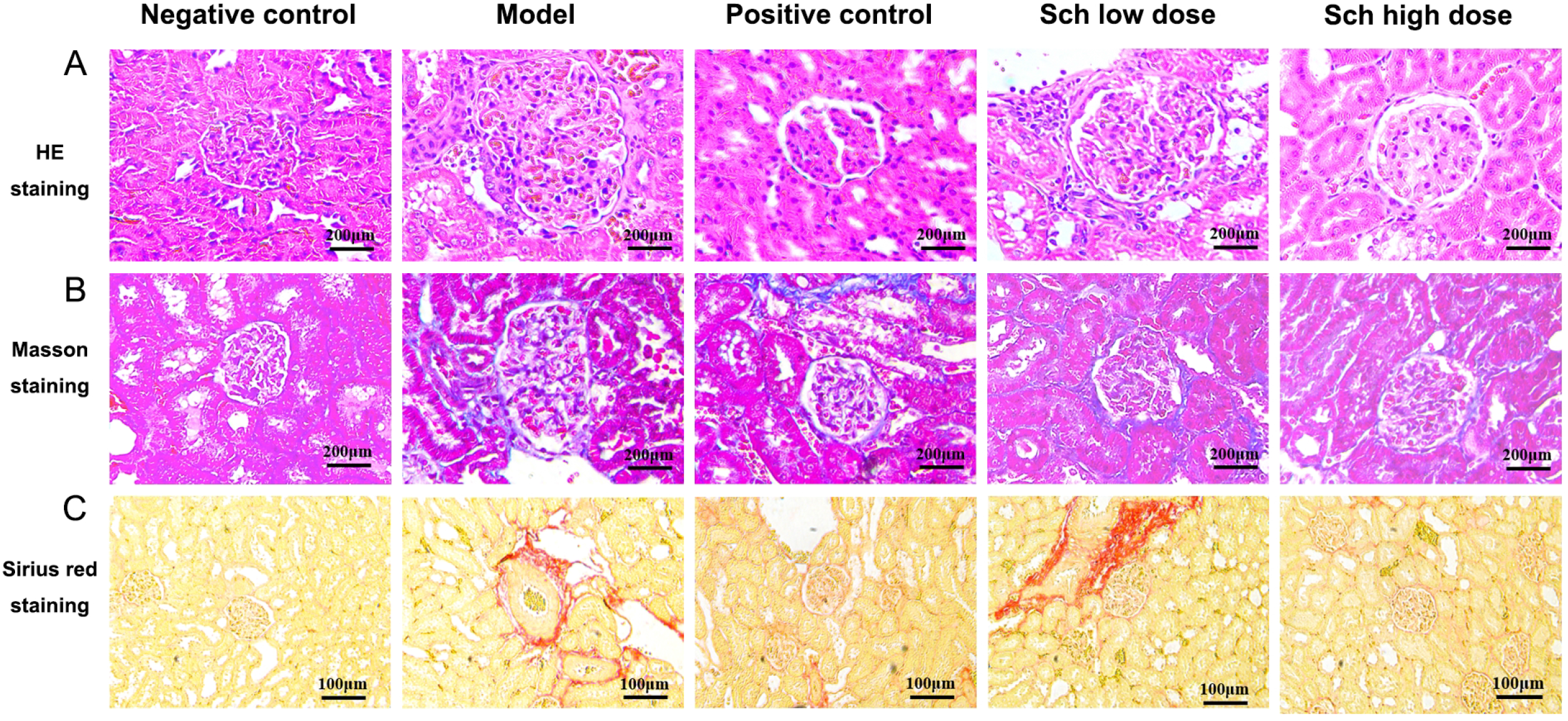
Sch treatment significantly improved the pathological changes of the kidneys in DN model rats. A. HE staining of a rat kidney. Bar = 200 μm. B. Masson staining of a rat kidney. Bar = 200 μm. C. Sirius red staining of a rat kidney. Bar = 100 μm.

### 3.2. Sch effects on oxidative stress and inflammatory factors in DN rats

The effects of Sch on oxidative stress in DN rats were further explored by measuring the SOD, MDA, and GSH-Px levels in the renal tissue homogenates of rats in each group. ELISA was used to determine the levels of pro-inflammatory factors IL-6, IL-1β, and TNF-α in the renal tissue homogenates of the rats in each group to evaluate the effect of Sch on the inflammatory response. When compared to the negative control group, the activities of SOD and GSH-Px in the renal tissue homogenate of the model group showed a significant decline (*P* < 0.001), while the level of MDA showed a significant surge (*P* < 0.001). Irbesartan and high dose Sch intervention promoted the activities of SOD (*P* < 0.001 and *P* < 0.001, respectively) (Figure 3A) and GSH-Px (*P* < 0.001 and *P* < 0.001, respectively) (Figure 3A and C) but diminished the level of MDA (*P* < 0.01 and *P* < 0.05, respectively) (Figure 3B) in renal cells. ELISA results showed that the serum levels of the pro-inflammatory factors IL-6, IL-1β, and TNF-α in the model group significantly increased (all *P* < 0.01) compared to the negative control group. In addition, irbesartan and high dose Sch intervention significantly reduced the serum levels of IL-6, IL-1β, and TNF-α in DN rats (Figure 3D). In summary, the results taken together substantiated that Sch produced a therapeutic effect in DN rats, especially at a high dose, and therefore, the high dose was selected for subsequent experiments.

**Figure 3.**
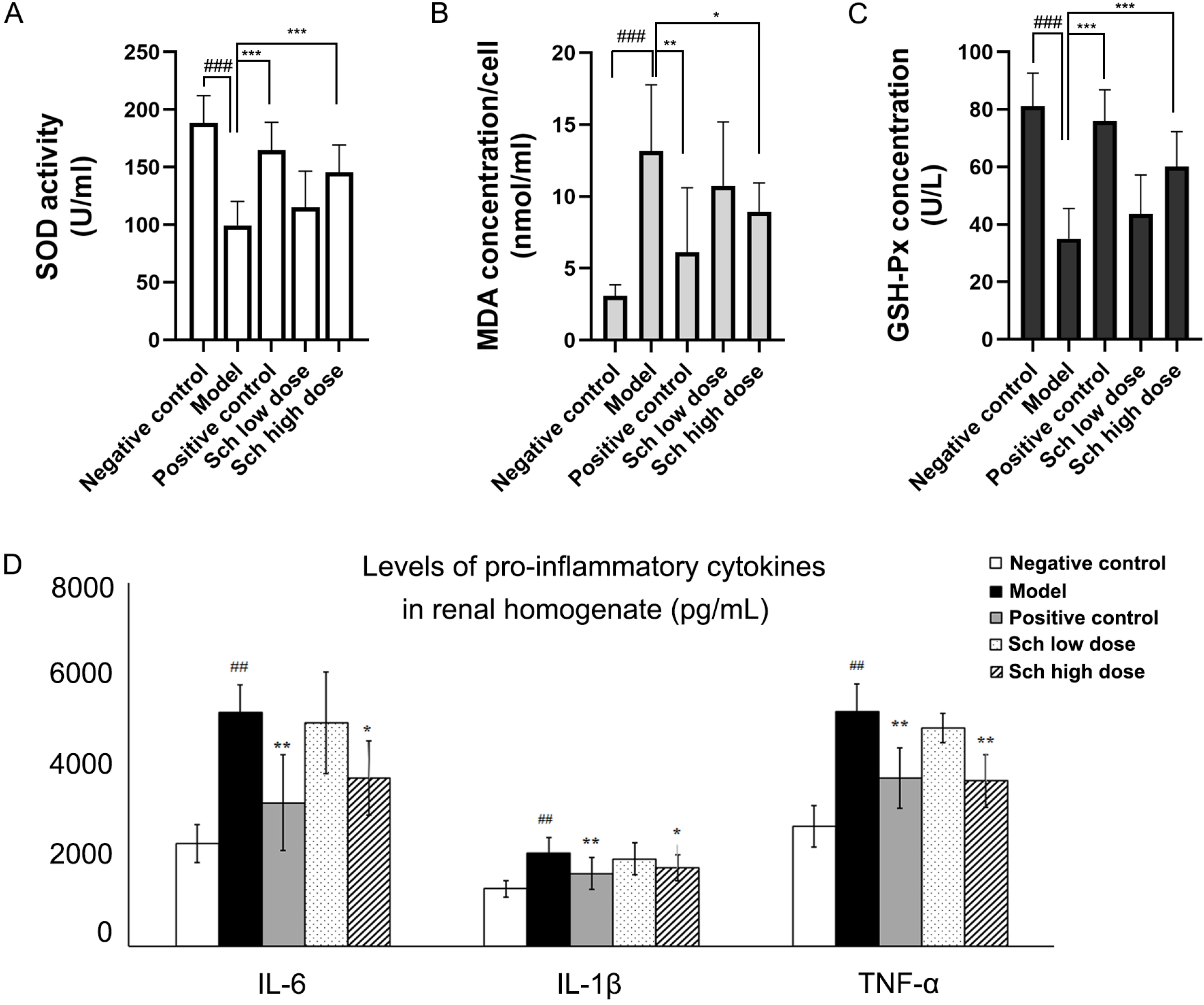
Sch showed anti-oxidative activity and inhibited the expression of pro-inflammatory cytokines in the serum of DN rats. A–C. SOD level (A) and MDA (B) and GSH-Px concentrations (C) in negative control and DN rats. D. Sch treatment inhibited the levels of pro-inflammatory cytokines in the serum. Negative control, model, positive control, Sch low dose, Sch high dose groups (n = 10 per group). Data are presented as the mean ± SD. ##: *P* < 0.01 as compared to the control group; ^*^: *P* < 0.05 as compared to the model group; ^**^: *P* < 0.01 as compared to the model group. ^***^: *P* < 0.001 as compared to the model group.

### 3.3. Network pharmacology analysis

With the aim of uncovering the effects of Sch on gene expression in the renal tissues of DN rats, network pharmacological prediction was used to elucidate the molecular mechanism of Sch in the treatment of DN, according to the Network Pharmacology Evaluation Method Guidance [39]. First, the ChEMBL and CTD databases were used to predict possible targets of Sch, and 34 target proteins were identified. According to the GeneCards and TCMSP databases, there were 3424 target proteins affected by DN disease. Eleven proteins, including XIAP associated factor 1 (XAF1), glutamic-pyruvic transaminase (GPT), glutamate-cysteine ligase catalytic subunit (GCLC), glutathione-disulfide reductase (GSR), FMS-related receptor tyrosine kinase 1 (FLT1), mitogen-activated protein kinase 1 (MAPK1), Crumbs cell polarity complex component 3 (Crb3), tetraspanin 4 (Tspan4), checkpoint kinase 1 (CHEK1), and mitogen-activated protein kinase 3 (MAPK3), and nuclear receptor subfamily 1 group I member 2 (NR1I2) were identified as targets of both Sch and DN disease by a Venn diagram (Figure 4A; Supplemental Table S1). GO analysis of the 11 proteins revealed that the top three biological processes in terms of enrichment were aging, apoptotic process, and protein phosphorylation (Figure 4B); the top three cell components in terms of enrichment were nucleus, mitochondrion, and nucleoplasm (Figure 4C); and the top three molecular functions in terms of enrichment were ATP binding, MAP kinase kinase activity, and MAP kinase activity (Figure 4D). KEGG pathway analysis was conducted with the purpose of discovering the enriched pathways of the 11 target genes. The results showed that the genes were enriched in the MAPK signaling pathway, PI3K-Akt signaling pathway, renal cell carcinoma, gap junction, endocrine resistance, and TNF signaling pathway (Figure 4E). Finally, a “bioactive-protein-pathways” interaction network was built to capture the relationship network between Sch, the target genes, and the related pathways (Figure 4F).

**Figure 4.**
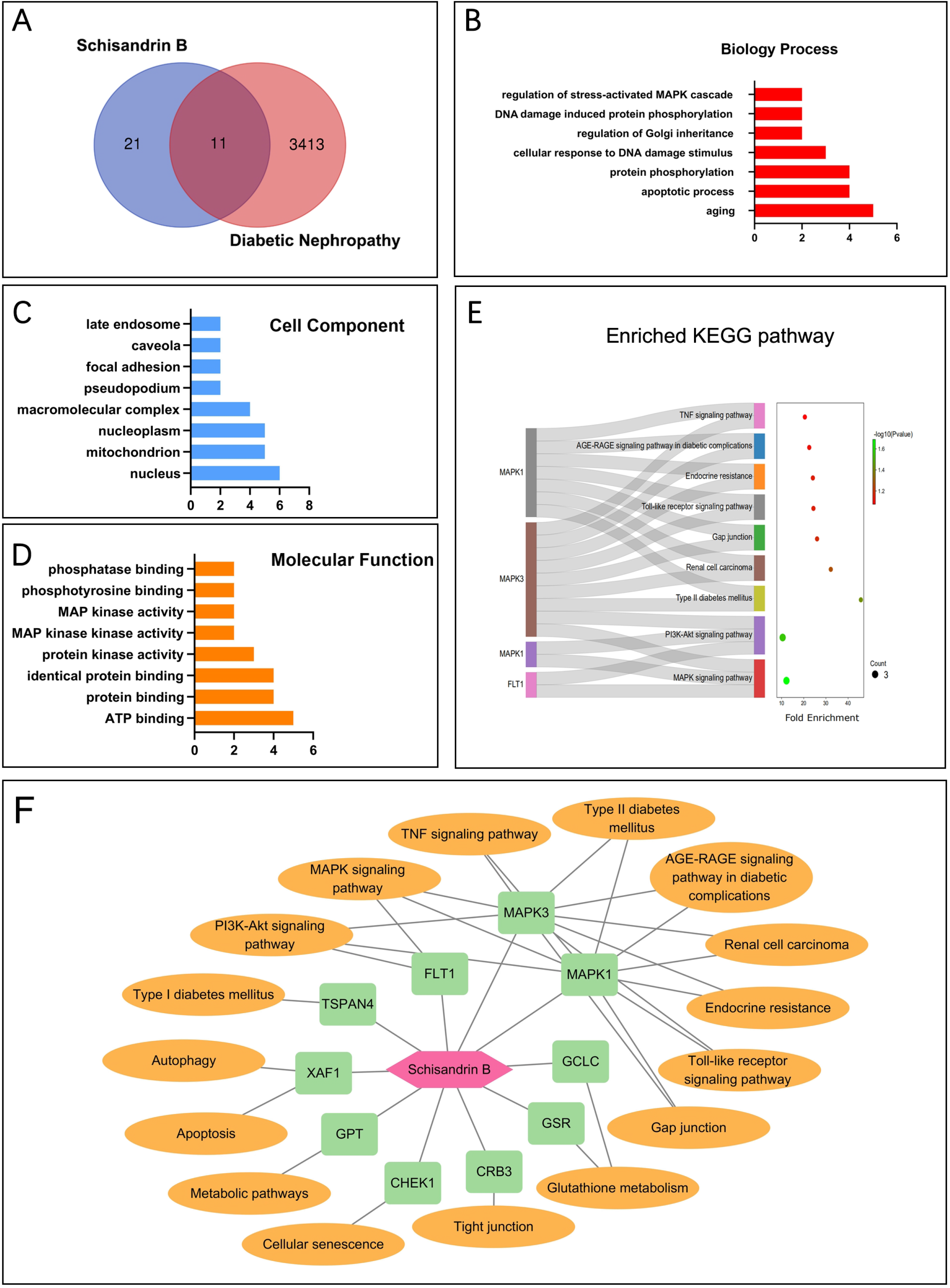
Network pharmacology analysis of intersecting genes between Sch targets and DN targets. A. Eleven intersecting genes were identified using Venn analysis. B–D. GO term analysis of the 11 genes, including biological process (B), cell component (C), and molecular function (D). E. KEGG analysis of the 11 genes. F. Bioactive-protein-pathway network of Sch, target genes, and the related pathways. The pink polygon represents Sch, the green rectangle represents the target genes, and the orange oval represents the related pathways.

### 3.4. Quantitative proteomics analysis

With the aim to detect protein expression changes caused by Sch in the kidneys of DN rats, proteomic sequencing was performed on the negative control group, model group, and Sch high dose group. The statistical results obtained of the spectrogram number, identified peptide number, and identified protein number are shown in Supplemental Figure 1. Using a screening standard of a fold change greater than 1.2 or lower than 0.83, as well as a *P* value < 0.05 *(t* test), the significantly differentially expressed proteins were effectively selected and separated into each group, indicating that the differentially expressed proteins could represent the effects of biological treatment on samples. The raw data of the quantitative proteomics with differentially expressed proteins marked with highlighting are presented in Supplemental Tables S2–S4.

First, we compared the overall differentially expressed proteins between the model vs. control group and between the Sch-treated vs. model group (Figure 5A and 5B). The differentially expressed protein, Xaf1, was found downregulated (0.80) in the model vs. control group and upregulated (1.24) in the Sch-treated vs. model group. Furthermore, five proteins were found upregulated in the model vs. control group and downregulated in the Sch-treated vs. model group: Crb3 (1.24; 0.80), Tspan4 (1.23; 0.77), WD repeat domain 45 (Wdr45) (1.36; 0.72), zinc finger protein 512 (Zfp512) (1.36; 0.73), and transmembrane and immunoglobulin domain containing 1 (Tmigd1) (1.26; 0.82). These proteins were considered as the possible targets of Sch in treating DN disease.

**Figure 5.**
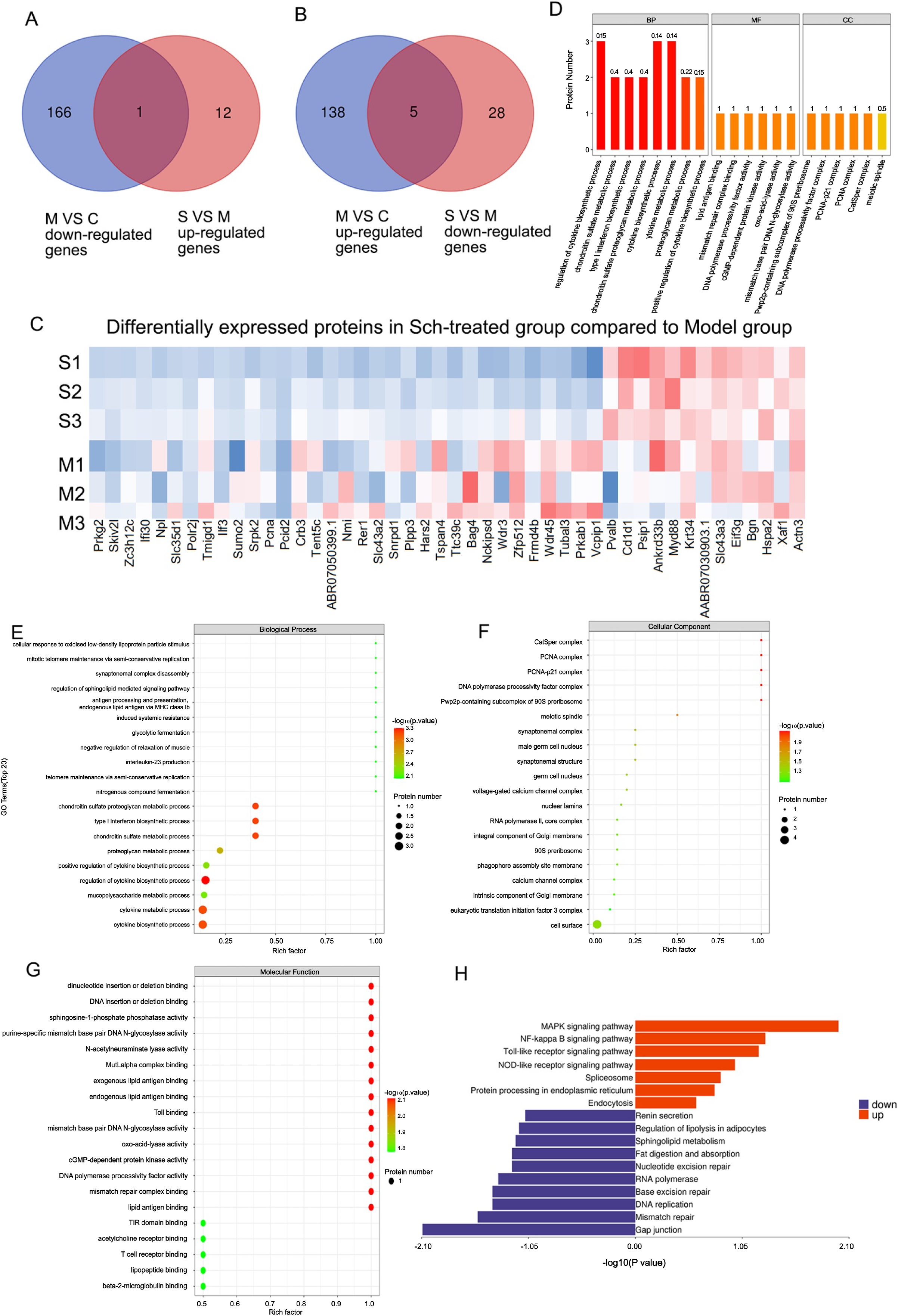
Quantitative proteomics analysis. A. Venn analysis of the upregulated proteins in the model vs. control group and downregulated proteins in the Sch-treated vs. model group. B. Venn analysis of the downregulated proteins in the model vs. control group and upregulated proteins in the Sch-treated vs. model group. C. Heatmap analysis of the differentially expressed proteins in the Sch-treated vs. model group. E–G. GO term analysis of the differentially expressed proteins in the Sch-treated vs. model group, including biological process (E), cell component (F), and molecular function (G). H. KEGG analysis of the differentially expressed proteins.

Next, we analyzed the differentially expressed proteins in the Sch-treated vs. model group (Figure 5C). GO term analysis was conducted to find the differentially expressed proteins. Fisher’s exact test was conducted to find the significant difference between the two and the functional categories of differentially expressed protein enrichment (*P* value < 0.05). The column diagram showed the general results of GO (Figure 5D), and the bubble diagram showed the enrichment of the three GO terms: biological process, cell component, and molecular function (Figure 5E–5G). The results showed that most genes were enriched in the following biological processes: cytokine biosynthetic process, cytokine metabolic process, mucopolysaccharide metabolic process, regulation of cytokine biosynthetic process, positive regulation of cytokine biosynthetic process, and proteoglycan metabolic process. Additionally, the following molecular functions underwent significant alterations: cGMP−dependent protein kinase activity, acetylcholine receptor binding, TIR domain binding, mismatch repair complex binding, and DNA polymerase processivity factor activity.

Protein coordination should be emphasized to gain a systematic and comprehensive understanding of a biological process, the mechanisms of disease occurrence, and the mechanisms of drug action. The Kyoto Encyclopedia of Genes and Genomes pathway (KEGG) database was utilized to analyze and annotate proteins. As shown in Figure 5H, among the differentially expressed protein pathways, the main enriched pathways were MAPK signaling, NF-κB signaling, Toll-like receptor signaling, NOD-like receptor signaling, gap junction, mismatch repair, regulation of lipolysis in adipocytes, and renin secretion (Figure 5H).

### 3.5. Intersected targets between network pharmacology prediction and proteomics

Finally, the intersected targets were analyzed using the Venn method. Three proteins, Crb3, Xaf1, and Tspan4, were identified in the network pharmacology prediction, and these were found to be differentially expressed in the Sch-treated vs. model group in the proteomics analysis (Figure 6A). The interacting network among the Sch-targets-pathway was then built with Cytoscape 3.7.0 (Figure 6B).

**Figure 6.**
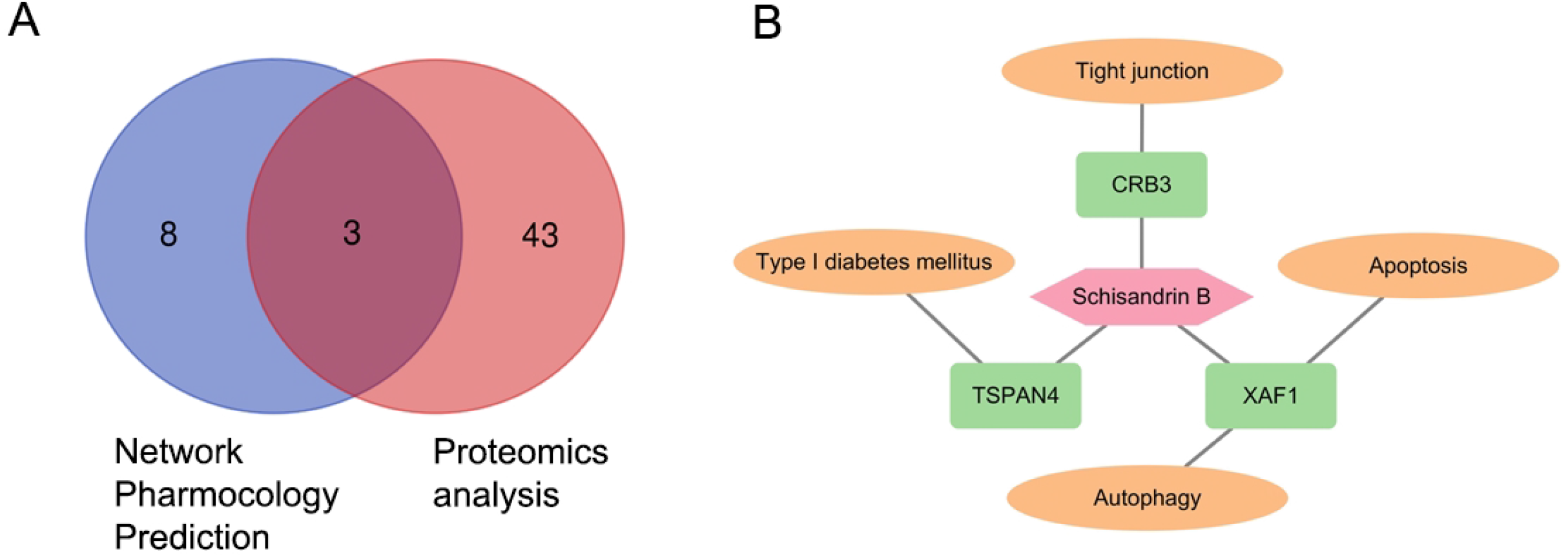
Three intersecting genes and the Sch-target-pathway network between the network pharmacology analysis and proteomics analysis. A. Three intersecting genes was identified using Venn analysis. B. Sch-target-pathway network. The pink rectangle represents Sch, the magenta triangles represent targets, and the blue circles represent pathways.

## 4. DISCUSSION

Studies on DN are regarded as crucial and fruitful, considering the high medical costs and cardiovascular deterioration of the population globally [40], and are needed to identify innovative treatment for diabetes [41-43]. Our work showed that Sch reduced blood glucose levels, improved biochemical indexes related to renal function, and alleviated renal damage in DN rats to different degrees, indicating that Sch has therapeutic effects on DN disease (Figure 1 and 2).

The treatment of DN varies according to different disease stages. Drug treatment, dialysis treatment, and renal transplantation are the main methods to treat DN [44], of which drug treatment is the most common. Hemodialysis can prolong the survival rate of patients, but infection, vascular calcification, injury, and other problems may occur after extended treatment [45]. Renal transplantation brings a heavy economic burden to patients, and it can neither prevent the recurrence of DN nor improve other diabetic complications. Therapeutic drugs include antihyperglycemic, antihypertensive, anti-inflammatory, antioxidant, and antifibrosis drugs, among which hyperglycemia control is the main means of treating DN. Dipeptidyl peptidase 4 (DPP-4) inhibitor is a new type 2 diabetes drug. It has been shown to increase the levels of endogenous glucagon-like peptide-1 (GLP-1) and glucose-dependent insulin-stimulating polypeptide (GIP). DPP-4 inhibitors promote pancreatic islet β cell insulin release and inhibit islet α cell glucagon secretion to increase insulin levels and reduce blood sugar. Matsui et al. reported that DPP-4 inhibitors blocked the level of (advanced glycation end products) AGEs and their receptors, reduced proteinuria, and reduced the degree of oxidative stress in a diabetic rat model [46]. Furthermore, Malek et al. showed that antagonists of the renin-angiotensin system (RAS) regulated renal hemodynamics, mainly through angiotensin II. RAS antagonists have the effects of lowering blood pressure, reducing urinary protein, and delaying the progression of DN disease [47]. Both Yoon et al. and Kang et al. found that angiotensin II induced selective changes in the glomerular basement membrane, increased proteinuria, produced oxidative stress through NADPH oxidase, and activated NF-κB, monocyte chemotactic protein-1 (MCP-1), and transforming growth factor-β (TGF-β) [48, 49]. These proteins were found to promote the generation of inflammation and renal fibrosis and to aggravate damage to the kidney [50]. However, studies have shown that there is a high risk of adverse reactions when such drugs are used [51]. Recently, Kimura et al. showed that sodium glucose cotransporter 2 (SGLT2) inhibitor inhibited the oxidative stress response by inhibiting the TLR signaling pathway, thereby alleviating renal damage in diabetic rats [52]. However, Wang et al. reported that although SGLT2 inhibitors reduced the incidence of hypoglycemia and acute renal injury in patients with type 2 diabetes, they carried a risk of urinary tract infections [53].

Nature is the most generous sponsor of drug research by providing abundant natural bioactive compounds, one of which is Sch. It is a non-enzymatic compound with low toxicity and low cost, and it is a multi-functional traditional medicine, and as such, it has an appealing drug prospect. Sch is a phenylcyclooctene lignan found at high levels in the fruit of *Schisandra chinensis*, a *Magnoliaceae* plant. It improves kidney function and has anti-inflammatory, antioxidant, antitumor, and other pharmacological effects. Lin et al. reported that Sch repressed the expression of Hsp27, downregulated inflammatory cytokines, and inhibited the occurrence of the inflammatory cascade reaction caused by p65 translocation to the nucleus [54]. Feng et al. showed that Sch was metabolized by the cytochrome P450 (CYP450s) enzyme to produce a carbonene active metabolite with the ability to modify peptides in human and mouse liver microsomes, which combined with Keap1 to activate the Nrf2 pathway to reduce liver damage [55]. Liu et al. showed that Sch activated NF-κB signal transduction, induced the expression of the antiapoptotic protein Survivin, and improved renal function damage caused by cisplatin, thus expanding the clinical application value of cisplatin [56]. Lai et al. found that Sch prevented the accumulation of reactive oxygen species by inhibiting oxidative stress, cell apoptosis, and autophagy and reduced the nephrotoxicity of the immunosuppressant cyclosporine A [57]. Whether Sch inhibits the pathogenesis of DN by regulating antioxidant pathways needs further verification.

Pessoa et al. showed that hyperglycemia caused chronic inflammation in vivo and promoted inflammatory cells to infiltrate into renal tissues, producing pro-inflammatory factors such as IL-1β, IL6, and TNF-α [58]. The release of these inflammatory factors further damaged the mesangial cells and epithelial cells, as well as the renal interstitium, and aggravated proteinuria. Toda et al. found that the levels of inflammatory factors were positively related to the severity of DN [59]. The high-dose Sch group had the most apparent effect on improving renal function, restoring pathological damage of renal tissues, and inhibiting inflammatory reactions in DN rats (Figure 3). These results showed that treatment with Sch had a significant anti-inflammatory effect on the kidneys of DN rats.

The proteomics analysis showed that proteins involved in MAPK signaling and the NF-κB signaling pathway were clustered in the Sch-treated DN rats. The MAPK signaling pathway regulates cell growth, differentiation, environmental stress adaptation, and inflammatory responses. Ran et al. reported that Sch prevented cartilage degeneration in a rat osteoarthritis model through inhibiting MAPK activation by reducing the expression level of p38, extracellular signal-regulated kinase, and c-Jun amino-terminal kinase phosphorylation [60]. Kim et al. also found that MAPK and TGF-β had different effects on the mesangial matrix compared to mesangial cells, resulting in glomerular hypertrophy in the progression of DN [61]. Liu et al. reported that metadherin induced podocyte apoptosis by activating the p38 MAPK-dependent pathway [62]. Furthermore, Sun et al. showed that the phosphorylation and acetylation levels of NF-κB p65 and signal transducer and activator of transcription 3 (Stat3) were upregulated during DN [63]. Whether Sch affects the epigenetic modification of NF-κB p65 and other proteins involved in MAPK and NF-κB signaling pathway needs further verification. For the prediction of direct interactions of small molecule compounds with target proteins, Almeida et al. isolated a mixture of flavonoids quercetin 3-O-methyl ether (1) and 6-C-methyl quercetin 3-O-methyl ether (2) from the ethyl acetate extract of aerial parts from Vellozia dasypus Seub And they studied the interaction of these two compounds with myeloperoxidase (MPO) by molecular docking [64]. Alves et al. used molecular modeling methods to predict the Raman spectroscopy, IR, and NMR data of piperine alkaloid crystals isolated from pepper (Piper nigrum L.) and compared the calculated data with the experimental data [65]. In future work, we will apply these methods for the prediction of this interaction, and the related validation work.

Tight junctions are cell-cell adhesions in epithelial and endothelial cellular sheets, acting as a primary barrier against the diffusion of solutes through the intracellular space (Figures 4 and 5). It creates a boundary between the apical and the basolateral plasma membrane domains and recruits various cytoskeletal and signaling molecules to their cytoplasmic surface [66]. As identified by our proteomics analysis, the following proteins might have roles in this process: tubulin alpha chain-like 3 (Tubal3), proliferating cell nuclear antigen (Pcna), and Crumbs protein homolog 3 (Crb3). We speculated that Sch may help the kidney create a boundary between the apical and the basolateral plasma membrane and recruit cytoskeletal and signaling molecules at the cytoplasmic surface of certain types of kidney cells.

Among the six possible targets of Sch in treating DN disease, Xaf1 was inhibited in the model vs. control group and induced in the Sch-treated vs. model group. Xaf1 is a negative regulator and its low expression level promoted autophagic cell death [67]. Xaf1 was downregulated in the model vs. control group and upregulated in the Sch-treated vs. model group (Supplemental Tables S2 and S4). This indicated that Sch may target Xaf1 and increase the expression of Xaf1 to reduce autophagy-related cell death during DN progression, to protect the kidneys of DN patients from damage. Tspan4 was induced in the model vs. control group and reduced in the Sch-treated vs. model group (Supplemental Tables S2 and S4). Ma et al. reported that Tspan4 was a novel histamine H4 receptor (H4R) interactor. H4R is a G protein-coupled receptor mainly expressed on immune cells. When phosphorylated by GPCR kinase, H4R activates G protein and recruits β-arrestins to induce cellular signals in response to agonist stimulation. H4R is considered a key drug target for various inflammatory disorders [68]. We hypothesized that Sch inhibited the inflammation response by inhibiting the expression of Tspan4 and interrupted the interaction between H4R and Tspan4. Zfp512 was reported to be associated with fat deposition and fat biosynthesis in pig muscles [69]. The inhibitory effect of Sch may reduce fat deposition and fat biosynthesis caused by diabetes. Tmigd1 has been shown to protect renal epithelial cells from injury induced by oxidative stress [70]. The suppression of Tmigd1 may be a feedback mechanism based on the induction of SOD or GSH-Px expression levels, or on the inhibition of MDA. Combined with the network pharmacology analysis, the intersected proteins, Crb3, Xaf1, and Tspan4, had the most potential to be Sch targets (Figure 6). The direct interaction between Sch and Crb3/Xaf1/Tspan4 will be verified in subsequent studies.

## 5. CONCLUSION

In summary, to investigate the possible mechanisms of Sch in treating DN, network pharmacology prediction and quantitative proteomics analysis showed that Sch treats DN by regulating the expression levels of Crb3, Xaf1 and Tspan4 to alleviate oxidative stress and inflammatory responses. These findings provide theoretical basis for elucidating the molecular mechanisms of Sch in treating DN.

## Supporting information

Affidavit of Approval of Animal Ethical and Welfare

Certificate of English Language Editing

## ETHICS APPROVAL AND CONSENT TO PARTICIPATE

The experiment was approved by the Ethics Committee of Tianjin University. Approval No. TJUE-2022-009

## HUMAN AND ANIMAL RIGHTS

No humans were used in this study. All animal research procedures were followed in accordance with the standards set forth in the eighth edition of Guide for the Care and Use of Laboratory Animals (published by the National Academy of Sciences, The National Academies Press, Washington, D.C.).

## CONSENT FOR PUBLICATION

Not applicable.

## AVAILABILITY OF DATA AND MATERIALS

Not applicable.

## ACKNOWLEDGEMENTS

We want to express our sincere gratitude to the laboratory partners who have patiently accompanied the entire writing and organizing process. We thank LetPub for its linguistic assistance during the preparation of this manuscript.

## AUTHOR CONTRIBUTIONS

Jianying Song: data analysis, visualization;

Bo Zhang: Funding acquisition;

Xudong Lyu: Writing-original draft;

Huiping Zhang: Data analysis;

Wenbo Cheng: Data analysis; writing-original draft;

Peiyuan Liu: Data analysis; writing-original draft;

Jun Kang: Conceptualization; writing-original draft; funding acquisition; supervision; writing-review and editing.

## SUPPLEMENTARY MATERIAL

Supplementary material is available on the publisher’s website along with the published article.

## Figure legends

**Scheme 1.**
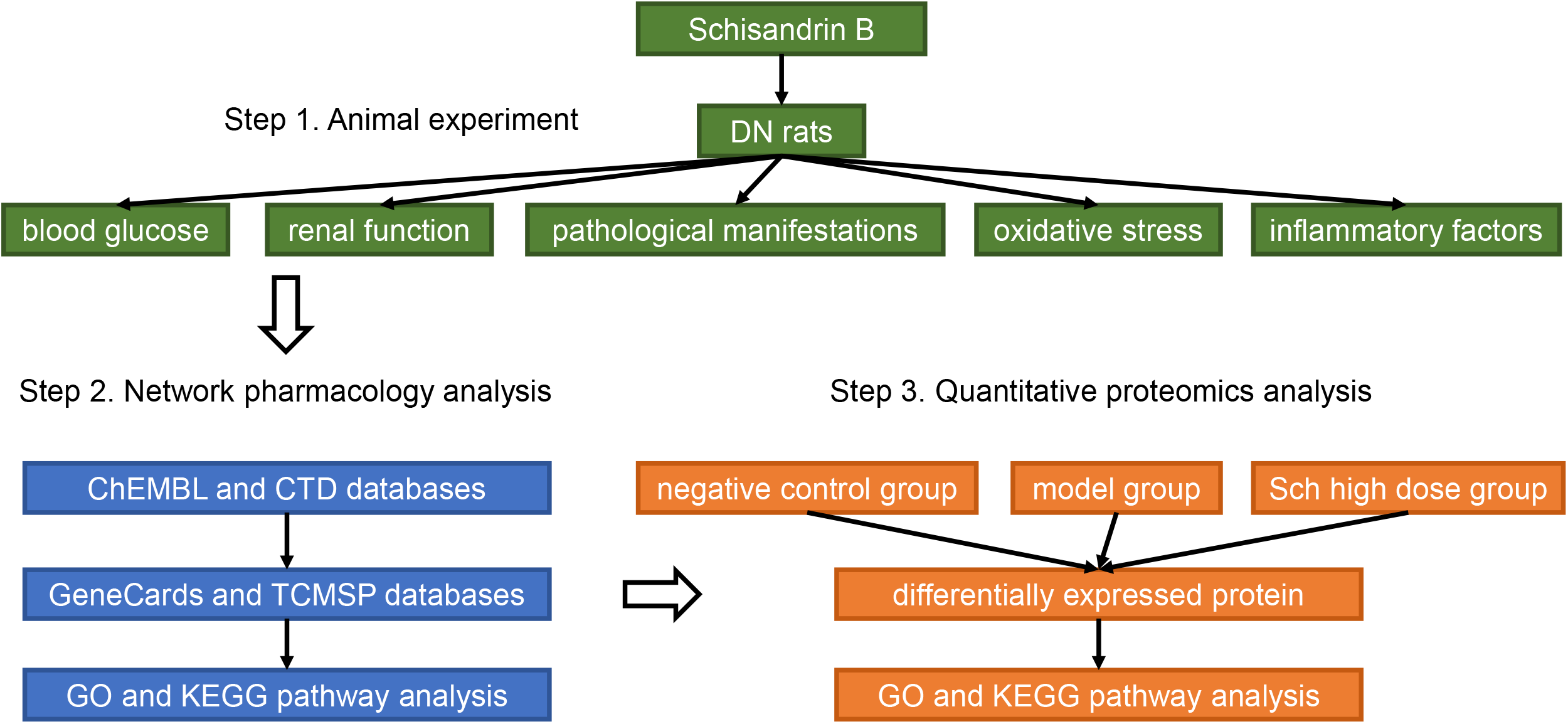
Schematic diagram of the experimental method flow.

## Supplemental materials

Supplemental Figure S1. Identified protein and peptide number from the quantitative proteomics.

Supplemental Table S1. Intersecting gene list between Schisandrin B target genes and diabetic nephropathy target genes.

Supplemental Table S2. Differentially expressed proteins in the model vs. control group.

Supplemental Table S3. Differentially expressed proteins in the Sch-treated vs. control group.

Supplemental Table S4. Differentially expressed proteins in the Sch-treated vs. model group.

## REFERENCES

[1] Goldberg, R.; Rubinstein, A.M.; Gil, N.; Hermano, E.; Li, J.P.; van der Vlag, J.; Atzmon, R.; Meirovitz, A.; Elkin, M. Role of heparanase-driven inflammatory cascade in pathogenesis of diabetic nephropathy. Diabetes, 2014, 63(12), 4302–4313. [https://doi.org/10.2337/db14-0001] [PMID: 25008182]

[2] Ni, W.J.; Tang, L.Q.; Wei, W. Research progress in signalling pathway in diabetic nephropathy. Diabetes Metab Res Rev, 2015, 31(3), 221–233. [https://doi.org/10.1002/dmrr.2568] [PMID: 24898554]

[3] Wang, H.; Zhang, H.; Chen, X.; Zhao, T.; Kong, Q.; Yan, M.; Zhang, B.; Sun, S.; Lan, H.Y.; Li, N.; Li, P. The decreased expression of electron transfer flavoprotein β is associated with tubular cell apoptosis in diabetic nephropathy. Int J Mol Med, 2016, 37(5), 1290–1298. [https://doi.org/10.3892/ijmm.2016.2533] [PMID: 27035869]

[4] Jin, H.; Piao, S.G.; Jin, J.Z.; Jin, Y.S.; Cui, Z.H.; Jin, H.F.; Zheng, H.L.; Li, J.J.; Jiang, Y.J.; Yang, C.W.; Li, C. Synergistic effects of leflunomide and benazepril in streptozotocin-induced diabetic nephropathy. Nephron Exp Nephrol, 2014, 126(3), 148–156. [https://doi.org/10.1159/000362556] [PMID: 24855017]

[5] Tuttle, K.R.; Bakris, G.L.; Toto, R.D.; McGill, J.B.; Hu, K.; Anderson, P.W. The effect of ruboxistaurin on nephropathy in type 2 diabetes. Diabetes Care, 2005, 28(11), 2686–2690. [https://doi.org/10.2337/diacare.28.11.2686] [PMID: 16249540]

[6] Hu, X.; Liu, W.; Yan, Y.; Liu, H.; Huang, Q.; Xiao, Y.; Gong, Z.; Du, J. Vitamin D protects against diabetic nephropathy: Evidence-based effectiveness and mechanism. Eur J Pharmacol, 2019, 845, 91–98. [https://doi.org/10.1016/j.ejphar.2018.09.037] [PMID: 30287151]

[7] Tuttle, K.R.; Brosius, F.C., 3rd; Adler, S.G.; Kretzler, M.; Mehta, R.L.; Tumlin, J.A.; Tanaka, Y.; Haneda, M.; Liu, J.; Silk, M.E.; Cardillo, T.E.; Duffin, K.L.; Haas, J.V.; Macias, W.L.; Nunes, F.P.; Janes, J.M. JAK1/JAK2 inhibition by baricitinib in diabetic kidney disease: results from a Phase 2 randomized controlled clinical trial. Nephrol Dial Transplant, 2018, 33(11), 1950–1959. [https://doi.org/10.1093/ndt/gfx377] [PMID: 29481660]

[8] Mora, C.; Navarro, J.F. Inflammation and diabetic nephropathy. Curr Diab Rep, 2006, 6(6), 463–468. [https://doi.org/10.1007/s11892-006-0080-1] [PMID: 17118230]

[9] Araújo, L.; Silva, M.; Silva, C.; Monteiro, M.; Pereira, L.; Rocha, L.P.; Corrêa, R.; Reis, M.A.; Machado, J.R. Cytokines and T Helper Cells in Diabetic Nephropathy Pathogenesis. 2016, 6(6), 230–246.

[10] Wada, J.; Makino, H. Inflammation and the pathogenesis of diabetic nephropathy. Clin Sci (Lond), 2013, 124(3), 139–152. [https://doi.org/10.1042/cs20120198] [PMID: 23075333]

[11] Lampropoulou, I.T.; Stanfigou, P.; Sarafidis, P.; Gouliovaki, A.; Giamalis, P.; Tsouchnikas, I.; Didangelos, T.; Papagianni, A. TNF-α pathway and T-cell immunity are activated early during the development of diabetic nephropathy in Type II Diabetes Mellitus. Clin Immunol, 2020, 215, 108423. [https://doi.org/10.1016/j.clim.2020.108423] [PMID: 32304735]

[12] Navarro-González, J.F.; Mora-Fernández, C. The role of inflammatory cytokines in diabetic nephropathy. J Am Soc Nephrol, 2008, 19(3), 433–442. [https://doi.org/10.1681/asn.2007091048] [PMID: 18256353]

[13] Börgeson, E.; Johnson, A.M.; Lee, Y.S.; Till, A.; Syed, G.H.; Ali-Shah, S.T.; Guiry, P.J.; Dalli, J.; Colas, R.A.; Serhan, C.N.; Sharma, K.; Godson, C. Lipoxin A4 Attenuates Obesity-Induced Adipose Inflammation and Associated Liver and Kidney Disease. Cell Metab, 2015, 22(1), 125–137. [https://doi.org/10.1016/j.cmet.2015.05.003] [PMID: 26052006]

[14] Liu, W.; Wu, Y.H.; Zhang, L.; Xue, B.; Wang, Y.; Liu, B.; Liu, X.Y.; Zuo, F.; Yang, X.Y.; Chen, F.Y.; Duan, R.; Cai, Y.; Zhang, B.; Ji, Y. MicroRNA-146a suppresses rheumatoid arthritis fibroblast-like synoviocytes proliferation and inflammatory responses by inhibiting the TLR4/NF-kB signaling. Oncotarget, 2018, 9(35), 23944–23959. [https://doi.org/10.18632/oncotarget.24050] [PMID: 29844864]

[15] Hu, S.; Zuo, H.; Qi, J.; Hu, Y.; Yu, B. Analysis of Effect of Schisandra in the Treatment of Myocardial Infarction Based on Three-Mode Gene Ontology Network. Front Pharmacol, 2019, 10, Article ID 232. [https://doi.org/10.3389/fphar.2019.00232] [PMID: 30949047]

[16] Thandavarayan, R.A.; Giridharan, V.V.; Arumugam, S.; Suzuki, K.; Ko, K.M.; Krishnamurthy, P.; Watanabe, K.; Konishi, T. Schisandrin B prevents doxorubicin induced cardiac dysfunction by modulation of DNA damage, oxidative stress and inflammation through inhibition of MAPK/p53 signaling. PLoS One, 2015, 10(3), e0119214. [https://doi.org/10.1371/journal.pone.0119214] [PMID: 25742619]

[17] Xu, Y.; Liu, Z.; Sun, J.; Pan, Q.; Sun, F.; Yan, Z.; Hu, X. Schisandrin B prevents doxorubicin-induced chronic cardiotoxicity and enhances its anticancer activity in vivo. PLoS One, 2011, 6(12), e28335. [https://doi.org/10.1371/journal.pone.0028335] [PMID: 22164272]

[18] Lee, T.H.; Jung, C.H.; Lee, D.H. Neuroprotective effects of Schisandrin B against transient focal cerebral ischemia in Sprague-Dawley rats. Food Chem Toxicol, 2012, 50(12), 4239–4245. [https://doi.org/10.1016/j.fct.2012.08.047] [PMID: 22960133]

[19] Wang, J.W.; Liang, F.Y.; Ouyang, X.S.; Li, P.B.; Pei, Z.; Su, W.W. Evaluation of neuroactive effects of ethanol extract of Schisandra chinensis, Schisandrin, and Schisandrin B and determination of underlying mechanisms by zebrafish behavioral profiling. Chin J Nat Med, 2018, 16(12), 916–925. [https://doi.org/10.1016/s1875-5364(18)30133-x] [PMID: 30595216]

[20] Yu, B.; Sheng, D.; Tan, Q. Determination of Schisandrin A and Schisandrin B in Traditional Chinese Medicine Preparation Huganpian Tablet by RP-HPLC. Chem Pharm Bull (Tokyo), 2019, 67(7), 713–716. [https://doi.org/10.1248/cpb.c18-00968] [PMID: 31006725]

[21] Qin, J.H.; Lin, J.R.; Ding, W.F.; Wu, W.H. Schisandrin B Improves the Renal Function of IgA Nephropathy Rats Through Inhibition of the NF-κB Signalling Pathway. Inflammation, 2019, 42(3), 884–894. [https://doi.org/10.1007/s10753-018-0943-z] [PMID: 30519926]

[22] Xu, J.; Lu, C.; Liu, Z.; Zhang, P.; Guo, H.; Wang, T. Schizandrin B protects LPS-induced sepsis via TLR4/NF-κB/MyD88 signaling pathway. Am J Transl Res, 2018, 10(4), 1155–1163. [PMID: 29736208]

[23] Li, M.; Jin, J.; Li, J.; Guan, C.W.; Wang, W.W.; Qiu, Y.W.; Huang, Z.Y. [Schisandrin B protects against nephrotoxicity induced by cisplatin in HK-2 cells via Nrf2-ARE activation]. Yao Xue Xue Bao, 2012, 47(11), 1434–1439. [PMID: 23387073]

[24] Mou, Z.; Feng, Z.; Xu, Z.; Zhuang, F.; Zheng, X.; Li, X.; Qian, J.; Liang, G. Schisandrin B alleviates diabetic nephropathy through suppressing excessive inflammation and oxidative stress. Biochem Biophys Res Commun, 2019, 508(1), 243–249. [https://doi.org/10.1016/j.bbrc.2018.11.128] [PMID: 30477745]

[25] Zhou, W.; Zhang, H.; Wang, X.; Kang, J.; Guo, W.; Zhou, L.; Liu, H.; Wang, M.; Jia, R.; Du, X.; Wang, W.; Zhang, B.; Li, S. Network pharmacology to unveil the mechanism of Moluodan in the treatment of chronic atrophic gastritis. Phytomedicine, 2022, 95, 153837. [https://doi.org/10.1016/j.phymed.2021.153837] [PMID: 34883416]

[26] 张 彦 琼 等. 网 络 药 理 学 与 中 医 药 现 代 研 究 的 若 干 进 展. 中 国 药 理 学 与 毒 理 学 杂 志, 2015, 29 (6):883–892.

[27] Li, S.; Zhang, B. Traditional Chinese medicine network pharmacology: theory, methodology and application. Chin J Nat Med, 2013, 11(2), 110–120. [https://doi.org/10.1016/s1875-5364(13)60037-0] [PMID: 23787177]

[28] Li, S. Mapping ancient remedies: Applying a network approach to traditional Chinese medicine. Science, 2015, 350(6262), S72–S74.

[29] Zhang, B.; Lu, C.; Bai, M.; He, X.; Tan, Y.; Bian, Y.; Xiao, C.; Zhang, G.; Lu, A.; Li, S. Tetramethylpyrazine identified by a network pharmacology approach ameliorates methotrexate-induced oxidative organ injury. J Ethnopharmacol, 2015, 175, 638–647. [https://doi.org/10.1016/j.jep.2015.09.034] [PMID: 26435225]

[30] Zhang, B.; Wang, X.; Li, Y.; Wu, M.; Wang, S.Y.; Li, S. Matrine Is Identified as a Novel Macropinocytosis Inducer by a Network Target Approach. Front Pharmacol, 2018, 9, 10. [https://doi.org/10.3389/fphar.2018.00010] [PMID: 29434546]

[31] Zhang, S.; Lai, X.; Wang, X.; Liu, G.; Wang, Z.; Cao, L.; Zhang, X.; Xiao, W.; Li, S. Deciphering the Pharmacological Mechanisms of Guizhi-Fuling Capsule on Primary Dysmenorrhea Through Network Pharmacology. Front Pharmacol, 2021, 12, 613104. [https://doi.org/10.3389/fphar.2021.613104] [PMID: 33746752]

[32] Guo, Y.; Bao, C.; Ma, D.; Cao, Y.; Li, Y.; Xie, Z.; Li, S. Network-Based Combinatorial CRISPR-Cas9 Screens Identify Synergistic Modules in Human Cells. ACS Synth Biol, 2019, 8(3), 482–490. [https://doi.org/10.1021/acssynbio.8b00237] [PMID: 30762338]

[33] Lei, Y.; Li, S.; Liu, Z.; Wan, F.; Tian, T.; Li, S.; Zhao, D.; Zeng, J. A deep-learning framework for multi-level peptide-protein interaction prediction. Nat Commun, 2021, 12(1), 5465. [https://doi.org/10.1038/s41467-021-25772-4] [PMID: 34526500]

[34] Zhou, W.; Lai, X.; Wang, X.; Yao, X.; Wang, W.; Li, S. Network pharmacology to explore the anti-inflammatory mechanism of Xuebijing in the treatment of sepsis. Phytomedicine, 2021, 85, 153543. [https://doi.org/10.1016/j.phymed.2021.153543] [PMID: 33799226]

[35] Zuo, J.; Wang, X.; Liu, Y.; Ye, J.; Liu, Q.; Li, Y.; Li, S. Integrating Network Pharmacology and Metabolomics Study on Anti-rheumatic Mechanisms and Antagonistic Effects Against Methotrexate-Induced Toxicity of Qing-Luo-Yin. Front Pharmacol, 2018, 9, 1472. [https://doi.org/10.3389/fphar.2018.01472] [PMID: 30618762]

[36] Azushima, K.; Gurley, S.B.; Coffman, T.M. Modelling diabetic nephropathy in mice. Nat Rev Nephrol, 2018, 14(1), 48–56. [https://doi.org/10.1038/nrneph.2017.142] [PMID: 29062142]

[37] Ge, J.; Miao, J.J.; Sun, X.Y.; Yu, J.Y. Huangkui capsule, an extract from Abelmoschus manihot (L.) medic, improves diabetic nephropathy via activating peroxisome proliferator-activated receptor (PPAR)-α/γ and attenuating endoplasmic reticulum stress in rats. J Ethnopharmacol, 2016, 189, 238–249. [https://doi.org/10.1016/j.jep.2016.05.033] [PMID: 27224243]

[38] Wiśniewski, J.R.; Zougman, A.; Nagaraj, N.; Mann, M. Universal sample preparation method for proteome analysis. Nat Methods, 2009, 6(5), 359–362. [https://doi.org/10.1038/nmeth.1322] [PMID: 19377485]

[39] Li, S.; Chen, Y.T.; Ding, Q.Y.; Dai, J.Y.; Duan, X.C.; Hu, Y.J.; Lai, X.X.; Liu, Q.F.; Niu, M.; Xiang, R.W. Network Pharmacology Evaluation Method Guidance-Draft. WJTCM, 2021, 7(1), 146–154.

[40] Meza Letelier, C.E.; San Martín Ojeda, C.A.; Ruiz Provoste, J.J.; Frugone Zaror, C.J. [Pathophysiology of diabetic nephropathy: a literature review]. Medwave, 2017, 17(1), e6839 [https://doi.org/10.5867/medwave.2017.01.6839] [PMID: 28112712]

[41] Gao, H.; Wu, H. Maslinic acid activates renal AMPK/SIRT1 signaling pathway and protects against diabetic nephropathy in mice. BMC Endocr Disord, 2022, 22(1), 25–35. [https://doi.org/10.1186/s12902-022-00935-6] [PMID: 35042497]

[42] Carmona, M.; Paco-Meza, L.M.; Ortega, R.; Cañadillas, S.; Caballero-Villarraso, J.; Blanco, A.; Herrera, C. Hypoxia preconditioning increases the ability of healthy but not diabetic rat-derived adipose stromal/stem cells (ASC) to improve histological lesions of streptozotocin-induced diabetic nephropathy. Pathol Res Pract, 2022, 230, Article ID 153756. [https://doi.org/10.1016/j.prp.2021.153756] [PMID: 35032832]

[43] Nagib, A.M.; Elsayed Matter, Y.; Gheith, O.A.; Refaie, A.F.; Othman, N.F.; Al-Otaibi, T. Diabetic Nephropathy Following Posttransplant Diabetes Mellitus. Exp Clin Transplant, 2019, 17(2), 138–146. [https://doi.org/10.6002/ect.2018.0157] [PMID: 30945628]

[44] Selby, N.M.; Taal, M.W. An updated overview of diabetic nephropathy: Diagnosis, prognosis, treatment goals and latest guidelines. Diabetes Obes Metab, 2020, 22(Suppl 1), 3–15. [https://doi.org/10.1111/dom.14007] [PMID: 32267079]

[45] Markell, M.S.; Friedman, E.A. Diabetic nephropathy. Management of the end-stage patient. Diabetes Care, 1992, 15(9), 1226–1238. [https://doi.org/10.2337/diacare.15.9.1226] [PMID: 1396019]

[46] Matsui, T.; Nakashima, S.; Nishino, Y.; Ojima, A.; Nakamura, N.; Arima, K.; Fukami, K.; Okuda, S.; Yamagishi, S. Dipeptidyl peptidase-4 deficiency protects against experimental diabetic nephropathy partly by blocking the advanced glycation end products-receptor axis. Lab Invest, 2015, 95(5), 525–533. [https://doi.org/10.1038/labinvest.2015.35] [PMID: 25730373]

[47] Malek, V.; Sharma, N.; Sankrityayan, H.; Gaikwad, A.B. Concurrent neprilysin inhibition and renin-angiotensin system modulations prevented diabetic nephropathy. Life Sci, 2019, 221, 159–167. [https://doi.org/10.1016/j.lfs.2019.02.027] [PMID: 30769114]

[48] Yoon, J.J.; Park, J.H.; Kim, H.J.; Jin, H.G.; Kim, H.Y.; Ahn, Y.M.; Kim, Y.C.; Lee, H.S.; Lee, Y.J.; Kang, D.G. Dianthus superbus Improves Glomerular Fibrosis and Renal Dysfunction in Diabetic Nephropathy Model. Nutrients, 2019, 11(3), 553–571. [https://doi.org/10.3390/nu11030553] [PMID: 30841605]

[49] Kang, J.S.; Lee, S.J.; Lee, J.H.; Kim, J.H.; Son, S.S.; Cha, S.K.; Lee, E.S.; Chung, C.H.; Lee, E.Y. Angiotensin II-mediated MYH9 downregulation causes structural and functional podocyte injury in diabetic kidney disease. Sci Rep, 2019, 9(1), 7679–7691. [https://doi.org/10.1038/s41598-019-44194-3] [PMID: 31118506]

[50] Ruggenenti, P.; Cravedi, P.; Remuzzi, G. The RAAS in the pathogenesis and treatment of diabetic nephropathy. Nat Rev Nephrol, 2010, 6(6), 319–330. [https://doi.org/10.1038/nrneph.2010.58] [PMID: 20440277]

[51] Harel, Z.; Gilbert, C.; Wald, R.; Bell, C.; Perl, J.; Juurlink, D.; Beyene, J.; Shah, P.S. The effect of combination treatment with aliskiren and blockers of the renin-angiotensin system on hyperkalaemia and acute kidney injury: systematic review and meta-analysis. BMJ, 2012, 344, Article ID e42. [https://doi.org/10.1136/bmj.e42] [PMID: 22232539]

[52] Kimura, Y.; Kuno, A.; Tanno, M.; Sato, T.; Ohno, K.; Shibata, S.; Nakata, K.; Sugawara, H.; Abe, K.; Igaki, Y.; Yano, T.; Miki, T.; Miura, T. Canagliflozin, a sodium-glucose cotransporter 2 inhibitor, normalizes renal susceptibility to type 1 cardiorenal syndrome through reduction of renal oxidative stress in diabetic rats. J Diabetes Investig, 2019, 10(4), 933–946. [https://doi.org/10.1111/jdi.13009] [PMID: 30663266]

[53] Wang, M.; Zhang, X.; Ni, T.; Wang, Y.; Wang, X.; Wu, Y.; Zhu, Z.; Li, Q. Comparison of New Oral Hypoglycemic Agents on Risk of Urinary Tract and Genital Infections in Type 2 Diabetes: A Network Meta-analysis. Adv Ther, 2021, 38(6), 2840–2853. [https://doi.org/10.1007/s12325-021-01759-x] [PMID: 33999339]

[54] Lin, Q.N.; Liu, Y.D.; Guo, S.E.; Zhou, R.; Huang, Q.; Zhang, Z.M.; Qin, X. Schisandrin B ameliorates high-glucose-induced vascular endothelial cells injury by regulating the Noxa/Hsp27/NF-κB signaling pathway. Biochem Cell Biol, 2019, 97(6), 681–692. [https://doi.org/10.1139/bcb-2018-0321] [PMID: 30817212]

[55] Feng, S.; Qiu, B.; Zou, L.; Liu, K.; Xu, X.; Zhu, H. Schisandrin B elicits the Keap1-Nrf2 defense system via carbene reactive metabolite which is less harmful to mice liver. Drug Des Devel Ther, 2018, 12, 4033–4046. [https://doi.org/10.2147/dddt.S176561] [PMID: 30568426]

[56] Liu, Q.; Song, J.; Li, H.; Dong, L.; Dai, S. Schizandrin B inhibits the cis-DDP-induced apoptosis of HK-2 cells by activating ERK/NF-κB signaling to regulate the expression of survivin. Int J Mol Med, 2018, 41(4), 2108–2116. [https://doi.org/10.3892/ijmm.2018.3409] [PMID: 29393335]

[57] Lai, Q.; Luo, Z.; Wu, C.; Lai, S.; Wei, H.; Li, T.; Wang, Q.; Yu, Y. Attenuation of cyclosporine A induced nephrotoxicity by schisandrin B through suppression of oxidative stress, apoptosis and autophagy. Int Immunopharmacol, 2017, 52, 15–23. [https://doi.org/10.1016/j.intimp.2017.08.019] [PMID: 28846887]

[58] Pessoa, E.A.; Convento, M.B.; Castino, B.; Leme, A.M.; de Oliveira, A.S.; Aragão, A.; Fernandes, S.M.; Carbonel, A.; Dezoti, C.; Vattimo, M.F.; Schor, N.; Borges, F.T. Beneficial Effects of Isoflavones in the Kidney of Obese Rats Are Mediated by PPAR-Gamma Expression. Nutrients, 2020, 12(6), 1624–1643. [https://doi.org/10.3390/nu12061624] [PMID: 32492810]

[59] Toda, N.; Mukoyama, M.; Yanagita, M.; Yokoi, H. CTGF in kidney fibrosis and glomerulonephritis. Inflamm Regen, 2018, 38, 14–21. [https://doi.org/10.1186/s41232-018-0070-0] [PMID: 30123390]

[60] Ran, J.; Ma, C.; Xu, K.; Xu, L.; He, Y.; Moqbel, S.A.A.; Hu, P.; Jiang, L.; Chen, W.; Bao, J.; Xiong, Y.; Wu, L. Schisandrin B ameliorated chondrocytes inflammation and osteoarthritis via suppression of NF-κB and MAPK signal pathways. Drug Des Devel Ther, 2018, 12, 1195–1204. [https://doi.org/10.2147/dddt.S162014] [PMID: 29785089]

[61] Kim, N.H. Podocyte hypertrophy in diabetic nephropathy. Nephrology (Carlton), 2005, 10, S14–16. [https://doi.org/10.1111/j.1440-1797.2005.00450.x] [PMID: 16174280]

[62] Liu, W.T.; Peng, F.F.; Li, H.Y.; Chen, X.W.; Gong, W.Q.; Chen, W.J.; Chen, Y.H.; Li, P.L.; Li, S.T.; Xu, Z.Z.; Long, H.B. Metadherin facilitates podocyte apoptosis in diabetic nephropathy. Cell Death Dis, 2016, 7(11), e2477. [https://doi.org/10.1038/cddis.2016.335] [PMID: 27882943]

[63] Sun, H.J.; Xiong, S.P.; Cao, X.; Cao, L.; Zhu, M.Y.; Wu, Z.Y.; Bian, J.S. Polysulfide-mediated sulfhydration of SIRT1 prevents diabetic nephropathy by suppressing phosphorylation and acetylation of p65 NF-κB and STAT3. Redox Biol, 2021, 38, 101813. [https://doi.org/10.1016/j.redox.2020.101813] [PMID: 33279869]

[64] Almeida, V.M.; Dias Ê, R.; Souza, B.C.; Cruz, J.N.; Santos, C.B.R.; Leite, F.H.A.; Queiroz, R.F.; Branco, A. Methoxylated flavonols from Vellozia dasypus Seub ethyl acetate active myeloperoxidase extract: in vitro and in silico assays. J Biomol Struct Dyn, 2022, 40(16), 7574–7583. [https://doi.org/10.1080/07391102.2021.1900916] [PMID: 33739225]

[65] Alves, F.S.; Rodrigues Do Rego, J.d.A.; Da Costa, M.L.; Lobato Da Silva, L.F.; Da Costa, R.A.; Cruz, J.N.; Brasil, D.D.S.B. Spectroscopic methods and in silico analyses using density functional theory to characterize and identify piperine alkaloid crystals isolated from pepper (Piper Nigrum L.). Journal of Biomolecular Structure and Dynamics, 2020, 38(9), 2792–2799. [https://doi.org/10.1080/07391102.2019.1639547]

[66] Tsukita, S. Tight Junctions. Encyclopedia of Biological Chemistry, 2013, 392-395. [https://doi.org/10.1016/B978-0-12-378630-2.00440-0]

[67] Sun, P.H.; Zhu, L.M.; Qiao, M.M.; Zhang, Y.P.; Jiang, S.H.; Wu, Y.L.; Tu, S.P. The XAF1 tumor suppressor induces autophagic cell death via upregulation of Beclin-1 and inhibition of Akt pathway. Cancer Lett, 2011, 310(2), 170–180. [https://doi.org/10.1016/j.canlet.2011.06.037] [PMID: 21788101]

[68] Ma, X.; Verweij, E.W.E.; Siderius, M.; Leurs, R.; Vischer, H.F. Identification of TSPAN4 as Novel Histamine H(4) Receptor Interactor. Biomolecules, 2021, 11(8), 1127–1142. [https://doi.org/10.3390/biom11081127] [PMID: 34439793]

[69] Ma, C.; Wang, W.; Wang, Y.; Sun, Y.; Kang, L.; Zhang, Q.; Jiang, Y. TMT-labeled quantitative proteomic analyses on the longissimus dorsi to identify the proteins underlying intramuscular fat content in pigs. J Proteomics, 2020, 213, 103630. [https://doi.org/10.1016/j.jprot.2019.103630] [PMID: 31881348]

[70] Wu, Y.; Li, E.; Wang, Z.; Shen, T.; Shen, C.; Liu, D.; Gao, Q.; Li, X.; Wei, G. TMIGD1 Inhibited Abdominal Adhesion Formation by Alleviating Oxidative Stress in the Mitochondria of Peritoneal Mesothelial Cells. Oxid Med Cell Longev, 2021, 9993704. [https://doi.org/10.1155/2021/9993704] [PMID: 34426761]

