## Supplemental for "Quantitative proteomics combined with network pharmacology analysis unveils the biological basis of Schisandrin B in treating diabetic nephropathy"

Song et al. Supplemental Figure S1

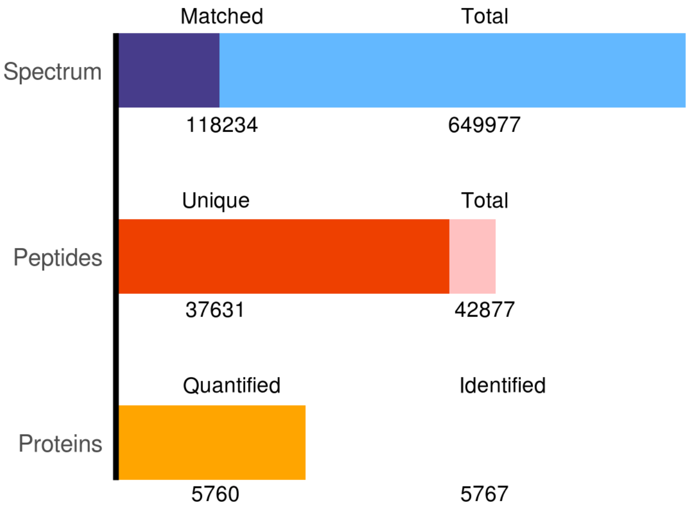

**Supplemental Table S1.** Intersecting gene list between Schisandrin B target genes and diabetic nephropathy target genes.

| Gene symbol |
|---|
| XAF1 |
| GPT |
| GCLC |
| GSR |
| FLT1 |
| MAPK1 |
| CRB3 |
| TSPAN4 |
| CHEK1 |
| MAPK3 |
| NR1I2 |

**Supplemental Table S2.** Differentially expressed proteins in the model vs. control group.

| Accession | Gene Name | M/C | t test p value |
|---|---|---|---|
| ENSRNOP00000041060 | Mt1m | 3.91 | 0.04 |
| ENSRNOP00000026122 | Hmgcs2 | 2.46 | 0.00 |
| ENSRNOP00000028664 | Mllt11 | 2.09 | 0.00 |
| ENSRNOP00000066577 | Gpx1 | 2.06 | 0.01 |
| ENSRNOP00000023835 | Akr1c14 | 1.74 | 0.01 |
| ENSRNOP00000073777 | Myo1g | 1.73 | 0.04 |
| ENSRNOP00000019795 | Sigmar1 | 1.69 | 0.01 |
| ENSRNOP00000030913 | Pck1 | 1.67 | 0.01 |
| ENSRNOP00000011784 | Eci1 | 1.66 | 0.00 |
| ENSRNOP00000015336 | Bhmt | 1.64 | 0.00 |
| ENSRNOP00000060140 | Acaa2 | 1.64 | 0.01 |
| ENSRNOP00000018622 | Rbp1 | 1.60 | 0.02 |
| ENSRNOP00000054044 | Naglt1 | 1.59 | 0.02 |
| ENSRNOP00000008613 | Slc5a8 | 1.58 | 0.02 |
| ENSRNOP00000043298 | Cyp24a1 | 1.58 | 0.00 |
| ENSRNOP00000002410 | Ehhadh | 1.57 | 0.03 |
| ENSRNOP00000005622 | S100g | 1.57 | 0.01 |
| ENSRNOP00000026878 | Hsd17b1 | 1.56 | 0.01 |
| ENSRNOP00000012759 | Pdk4 | 1.56 | 0.00 |
| ENSRNOP00000024622 | LOC501233 | 1.54 | 0.00 |
| ENSRNOP00000021694 | Vnn1 | 1.53 | 0.00 |
| ENSRNOP00000017712 | Nudt2 | 1.53 | 0.00 |
| ENSRNOP00000011945 | Gpnmb | 1.51 | 0.01 |
| ENSRNOP00000072741 | Map3k4 | 1.49 | 0.03 |
| ENSRNOP00000065846 | Wwc3 | 1.49 | 0.00 |
| ENSRNOP00000055007 | Tsc22d1 | 1.49 | 0.00 |
| ENSRNOP00000018737 | Gls2 | 1.49 | 0.00 |
| ENSRNOP00000005641 | Pdk2 | 1.45 | 0.01 |
| ENSRNOP00000027537 | Ech1 | 1.45 | 0.00 |
| ENSRNOP00000066155 | Amt | 1.45 | 0.01 |
| ENSRNOP00000073662 | Pigr | 1.44 | 0.00 |
| ENSRNOP00000074206 | Ephx1 | 1.44 | 0.03 |
| ENSRNOP00000027234 | Slc16a1 | 1.43 | 0.01 |
| ENSRNOP00000068322 | Tmlhe | 1.42 | 0.01 |
| ENSRNOP00000054540 | Ttc39a | 1.42 | 0.00 |
| ENSRNOP00000041451 | Hba-a3 | 1.41 | 0.03 |
| ENSRNOP00000028098 | Rnls | 1.40 | 0.01 |
| ENSRNOP00000014213 | Acot7 | 1.39 | 0.01 |
| ENSRNOP00000023739 | Akr1c3 | 1.39 | 0.04 |
| ENSRNOP00000060590 | Naga | 1.39 | 0.01 |
| ENSRNOP00000009777 | Baat | 1.38 | 0.01 |
| ENSRNOP00000075331 | Ddr1 | 1.38 | 0.00 |
| ENSRNOP00000050691 | Acaa1a | 1.38 | 0.04 |
| ENSRNOP00000008337 | Aco1 | 1.38 | 0.01 |
| ENSRNOP00000002712 | Ugt2b35 | 1.38 | 0.02 |
| ENSRNOP00000043148 | Comt | 1.38 | 0.01 |
| ENSRNOP00000022713 | Itga8 | 1.37 | 0.02 |

| | | | |
|---|---|---|---|
| ENSRNOP00000009462 | Adhfe1 | 1.37 | 0.02 |
| ENSRNOP00000004463 | Gulp1 | 1.37 | 0.02 |
| ENSRNOP00000072070 | Evl | 1.36 | 0.04 |
| ENSRNOP00000072771 | Trim14 | 1.36 | 0.00 |
| ENSRNOP00000058950 | Smim22 | 1.36 | 0.01 |
| ENSRNOP00000013369 | Wdr45 | 1.36 | 0.00 |
| ENSRNOP00000024000 | Aldh1a1 | 1.36 | 0.05 |
| ENSRNOP00000006658 | Zfp512 | 1.36 | 0.00 |
| ENSRNOP00000065206 | Apoa1 | 1.35 | 0.04 |
| ENSRNOP00000068639 | Gsta3 | 1.34 | 0.01 |
| ENSRNOP00000016784 | Trio | 1.34 | 0.02 |
| ENSRNOP00000038073 | Hadha | 1.34 | 0.01 |
| ENSRNOP00000041926 | Slc5a12 | 1.34 | 0.03 |
| ENSRNOP00000039445 | Espn | 1.34 | 0.01 |
| ENSRNOP00000005551 | Adipor1 | 1.33 | 0.01 |
| ENSRNOP00000055038 | Gstt3 | 1.33 | 0.00 |
| ENSRNOP00000016445 | Hddc3 | 1.33 | 0.00 |
| ENSRNOP00000007104 | Cnrip1 | 1.33 | 0.00 |
| ENSRNOP00000004673 | Acsf2 | 1.33 | 0.04 |
| ENSRNOP00000070446 | Vdr | 1.33 | 0.00 |
| ENSRNOP00000057148 | Adtrp | 1.31 | 0.00 |
| ENSRNOP00000014637 | Hadhb | 1.31 | 0.00 |
| ENSRNOP00000016954 | Cpt2 | 1.31 | 0.01 |
| ENSRNOP00000001365 | Ccz1b | 1.31 | 0.00 |
| ENSRNOP00000065713 | Akr1b8 | 1.30 | 0.03 |
| ENSRNOP00000021284 | Etfrf1 | 1.30 | 0.01 |
| ENSRNOP00000008254 | Lztfl1 | 1.30 | 0.02 |
| ENSRNOP00000075572 | Gsta4 | 1.30 | 0.01 |
| ENSRNOP00000066523 | Ctsz | 1.30 | 0.00 |
| ENSRNOP00000064322 | Dhrs7l1 | 1.30 | 0.01 |
| ENSRNOP00000023430 | Cep192 | 1.29 | 0.00 |
| ENSRNOP00000051538 | Acox1 | 1.29 | 0.04 |
| ENSRNOP00000044955 | Paqr9 | 1.29 | 0.00 |
| ENSRNOP00000040341 | Apon | 1.29 | 0.01 |
| ENSRNOP00000025800 | Gstm3l | 1.29 | 0.00 |
| ENSRNOP00000064884 | Ces2c | 1.29 | 0.00 |
| ENSRNOP00000010760 | Mettl7b | 1.29 | 0.00 |
| ENSRNOP00000061162 | RGD1565355 | 1.28 | 0.00 |
| ENSRNOP00000012455 | Fuca1 | 1.28 | 0.01 |
| ENSRNOP00000024659 | Slc27a1 | 1.28 | 0.00 |
| ENSRNOP00000010811 | Ndrg1 | 1.28 | 0.03 |
| ENSRNOP00000055608 | Acsl6 | 1.28 | 0.03 |
| ENSRNOP00000004947 | Mrpl27 | 1.27 | 0.01 |
| ENSRNOP00000065593 | Ces2c | 1.27 | 0.02 |
| ENSRNOP00000025859 | Myh7b | 1.27 | 0.02 |
| ENSRNOP00000053510 | Gp1ba | 1.27 | 0.01 |
| ENSRNOP00000013703 | Enpp5 | 1.27 | 0.00 |
| ENSRNOP00000040878 | LOC108351137 | 1.27 | 0.04 |
| ENSRNOP00000005129 | Tmigd1 | 1.27 | 0.03 |
| ENSRNOP00000024196 | Gpld1 | 1.26 | 0.00 |

| | | | |
|---|---|---|---|
| ENSRNOP00000006272 | Cdcp1 | 1.26 | 0.04 |
| ENSRNOP00000009627 | Acss1 | 1.26 | 0.03 |
| ENSRNOP00000026748 | Itgam | 1.26 | 0.02 |
| ENSRNOP00000009464 | Thumpd3 | 1.25 | 0.01 |
| ENSRNOP00000073800 | Apmap | 1.25 | 0.00 |
| ENSRNOP00000065600 | Ahsa1 | 1.25 | 0.00 |
| ENSRNOP00000023437 | Nudt6 | 1.25 | 0.03 |
| ENSRNOP00000026038 | Alox15 | 1.25 | 0.01 |
| ENSRNOP00000007927 | Rpia | 1.24 | 0.04 |
| ENSRNOP00000069820 | LOC103689961 | 1.24 | 0.02 |
| ENSRNOP00000064264 | Vav1 | 1.24 | 0.05 |
| ENSRNOP00000064452 | Crb3 | 1.24 | 0.01 |
| ENSRNOP00000038628 | Dgka | 1.24 | 0.02 |
| ENSRNOP00000012888 | Cyp4a1 | 1.24 | 0.00 |
| ENSRNOP00000000047 | Slc26a1 | 1.24 | 0.01 |
| ENSRNOP00000000203 | Csf2rb | 1.24 | 0.00 |
| ENSRNOP00000028732 | Ctss | 1.24 | 0.01 |
| ENSRNOP00000060618 | Rabl6 | 1.24 | 0.00 |
| ENSRNOP00000024093 | Aldh1a7 | 1.24 | 0.03 |
| ENSRNOP00000017468 | Ldha | 1.24 | 0.00 |
| ENSRNOP00000064289 | Vps26b | 1.23 | 0.00 |
| ENSRNOP00000020419 | Spryd7 | 1.23 | 0.01 |
| ENSRNOP00000052126 | Cep250 | 1.23 | 0.00 |
| ENSRNOP00000066775 | Tspan4 | 1.23 | 0.01 |
| ENSRNOP00000072623 | LOC681544 | 1.23 | 0.00 |
| ENSRNOP00000020374 | Tpd52l2 | 1.23 | 0.03 |
| ENSRNOP00000016176 | Tmem82 | 1.23 | 0.00 |
| ENSRNOP00000028194 | Dhcr7 | 1.23 | 0.00 |
| ENSRNOP00000031941 | Dsg4 | 1.23 | 0.03 |
| ENSRNOP00000003320 | Niban1 | 1.22 | 0.02 |
| ENSRNOP00000001466 | Foxk1 | 1.22 | 0.01 |
| ENSRNOP00000072412 | Mrpl14 | 1.22 | 0.02 |
| ENSRNOP00000071112 | Gstt2 | 1.22 | 0.03 |
| ENSRNOP00000075534 | Acot12 | 1.22 | 0.01 |
| ENSRNOP00000001638 | Pttg1ip | 1.22 | 0.03 |
| ENSRNOP00000069980 | Pacsin1 | 1.21 | 0.05 |
| ENSRNOP00000072048 | Metap1d | 1.21 | 0.01 |
| ENSRNOP00000030271 | Mgll | 1.21 | 0.00 |
| ENSRNOP00000001960 | Tmem120a | 1.21 | 0.05 |
| ENSRNOP00000031836 | Yod1 | 1.21 | 0.00 |
| ENSRNOP00000013607 | Slc27a2 | 1.21 | 0.02 |
| ENSRNOP00000074544 | Cyp2a1 | 1.21 | 0.03 |
| ENSRNOP00000049160 | Stradb | 1.21 | 0.00 |
| ENSRNOP00000042132 | Acox1 | 1.21 | 0.01 |
| ENSRNOP00000016148 | Rhpn2 | 1.20 | 0.00 |
| ENSRNOP00000025147 | Impa2 | 1.20 | 0.01 |
| ENSRNOP00000068477 | Unc5cl | 1.20 | 0.00 |
| ENSRNOP00000010079 | Asns | 1.20 | 0.01 |
| ENSRNOP00000051747 | Lsp1 | 0.83 | 0.02 |
| ENSRNOP00000003282 | Gpc4 | 0.83 | 0.00 |

| | | | |
|---|---|---|---|
| ENSRNOP00000018170 | Mat2a | 0.83 | 0.00 |
| ENSRNOP00000014496 | RGD1309534 | 0.83 | 0.00 |
| ENSRNOP00000003092 | Casr | 0.83 | 0.03 |
| ENSRNOP00000013058 | Nos3 | 0.83 | 0.00 |
| ENSRNOP00000025904 | Stom | 0.82 | 0.02 |
| ENSRNOP00000017139 | Pcbp4 | 0.82 | 0.00 |
| ENSRNOP00000074126 | LOC100365958 | 0.82 | 0.00 |
| ENSRNOP00000071291 | Slc5a6 | 0.82 | 0.00 |
| ENSRNOP00000012608 | Fkbp1a | 0.82 | 0.01 |
| ENSRNOP00000027220 | Scly | 0.82 | 0.00 |
| ENSRNOP00000024160 | Akr7a3 | 0.82 | 0.02 |
| ENSRNOP00000001389 | Slc25a30 | 0.82 | 0.01 |
| ENSRNOP00000005844 | Pah | 0.82 | 0.01 |
| ENSRNOP00000015888 | Vill | 0.82 | 0.02 |
| ENSRNOP00000073488 | Otud7b | 0.82 | 0.02 |
| ENSRNOP00000028474 | Bcat2 | 0.82 | 0.03 |
| ENSRNOP00000002579 | Ranbp1 | 0.82 | 0.04 |
| ENSRNOP00000011220 | Fut8 | 0.82 | 0.01 |
| ENSRNOP00000012300 | Fut11 | 0.82 | 0.04 |
| ENSRNOP00000064116 | Klhdc7a | 0.82 | 0.00 |
| ENSRNOP00000049903 | Ggct | 0.82 | 0.01 |
| ENSRNOP00000066885 | C3 | 0.82 | 0.04 |
| ENSRNOP00000066879 | LOC498122 | 0.81 | 0.05 |
| ENSRNOP00000066553 | LOC684270 | 0.81 | 0.00 |
| ENSRNOP00000023080 | Qprt | 0.81 | 0.02 |
| ENSRNOP00000068285 | AABR07048475.1 | 0.81 | 0.01 |
| ENSRNOP00000037089 | Slco4c1 | 0.81 | 0.02 |
| ENSRNOP00000026597 | Pgpep1 | 0.81 | 0.02 |
| ENSRNOP00000071311 | Folr1 | 0.81 | 0.01 |
| ENSRNOP00000048676 | Atp11c | 0.81 | 0.00 |
| ENSRNOP00000058050 | Rbm12b | 0.81 | 0.01 |
| ENSRNOP00000036519 | Dhrs7l1 | 0.81 | 0.03 |
| ENSRNOP00000043021 | AABR07044570.1 | 0.81 | 0.02 |
| ENSRNOP00000034652 | Cndp1 | 0.81 | 0.01 |
| ENSRNOP00000018003 | Hykk | 0.81 | 0.01 |
| ENSRNOP00000063373 | Ccdc88a | 0.81 | 0.04 |
| ENSRNOP00000068434 | Gosr1 | 0.81 | 0.03 |
| ENSRNOP00000006597 | Slc25a35 | 0.81 | 0.00 |
| ENSRNOP00000073834 | Dennd1a | 0.81 | 0.02 |
| ENSRNOP00000060972 | Hist3h2ba | 0.80 | 0.04 |
| ENSRNOP00000065710 | AABR07051551.2 | 0.80 | 0.01 |
| ENSRNOP00000067600 | Rab29 | 0.80 | 0.03 |
| ENSRNOP00000073116 | Asah2 | 0.80 | 0.01 |
| ENSRNOP00000071707 | Slc39a7 | 0.80 | 0.01 |
| ENSRNOP00000049378 | Pcid2 | 0.80 | 0.00 |
| ENSRNOP00000043362 | Mup4 | 0.80 | 0.03 |
| ENSRNOP00000053391 | Xaf1 | 0.80 | 0.05 |
| ENSRNOP00000039536 | Ighm | 0.80 | 0.02 |
| ENSRNOP00000048768 | Myl10 | 0.80 | 0.02 |
| ENSRNOP00000018837 | Wwp2 | 0.80 | 0.02 |

| | | | |
|---|---|---|---|
| ENSRNOP00000002540 | Sdf2l1 | 0.80 | 0.02 |
| ENSRNOP00000019767 | Fahd1 | 0.79 | 0.00 |
| ENSRNOP00000072092 | Dpp4 | 0.79 | 0.00 |
| ENSRNOP00000023116 | Slc22a2 | 0.79 | 0.02 |
| ENSRNOP00000041933 | AABR07051532.2 | 0.79 | 0.04 |
| ENSRNOP00000059987 | Acss2 | 0.79 | 0.03 |
| ENSRNOP00000003189 | Slc15a2 | 0.79 | 0.02 |
| ENSRNOP00000006954 | AABR07051626.2 | 0.79 | 0.00 |
| ENSRNOP00000022113 | Ttr | 0.79 | 0.00 |
| ENSRNOP00000040237 | Slco1a4 | 0.79 | 0.00 |
| ENSRNOP00000023274 | Cdk11b | 0.79 | 0.00 |
| ENSRNOP00000011384 | Mthfr | 0.78 | 0.01 |
| ENSRNOP00000068163 | Ly6e | 0.78 | 0.05 |
| ENSRNOP00000075613 | Arhgap24 | 0.78 | 0.00 |
| ENSRNOP00000058914 | LOC103691744 | 0.78 | 0.02 |
| ENSRNOP00000039968 | Cbs | 0.78 | 0.00 |
| ENSRNOP00000031764 | Slc34a1 | 0.78 | 0.00 |
| ENSRNOP00000038185 | Slc13a2 | 0.78 | 0.00 |
| ENSRNOP00000061340 | Slc2a1 | 0.78 | 0.01 |
| ENSRNOP00000020192 | Hagh | 0.77 | 0.00 |
| ENSRNOP00000029808 | Fdft1 | 0.77 | 0.00 |
| ENSRNOP00000000707 | Marcks | 0.77 | 0.05 |
| ENSRNOP00000052452 | Smim26 | 0.77 | 0.03 |
| ENSRNOP00000011941 | Miox | 0.77 | 0.01 |
| ENSRNOP00000004298 | Anks3 | 0.77 | 0.00 |
| ENSRNOP00000073568 | Arg2 | 0.77 | 0.01 |
| ENSRNOP00000072454 | AABR07065714.1 | 0.77 | 0.01 |
| ENSRNOP00000042580 | AABR07065768.1 | 0.77 | 0.02 |
| ENSRNOP00000015756 | Rpl22l1 | 0.77 | 0.02 |
| ENSRNOP00000015843 | Blvra | 0.77 | 0.00 |
| ENSRNOP00000045992 | Myl12b | 0.76 | 0.00 |
| ENSRNOP00000021027 | Umod | 0.76 | 0.03 |
| ENSRNOP00000070467 | Slc5a9 | 0.76 | 0.00 |
| ENSRNOP00000063111 | Wasf2 | 0.76 | 0.00 |
| ENSRNOP00000076230 | Mrps17 | 0.76 | 0.01 |
| ENSRNOP00000016912 | Mgam | 0.76 | 0.00 |
| ENSRNOP00000005440 | Tcaim | 0.75 | 0.00 |
| ENSRNOP00000055169 | Akr1c1 | 0.75 | 0.00 |
| ENSRNOP00000072904 | Arhgef1 | 0.75 | 0.02 |
| ENSRNOP00000007615 | Galnt3 | 0.75 | 0.01 |
| ENSRNOP00000015692 | Aqp1 | 0.75 | 0.02 |
| ENSRNOP00000013520 | Cldn10 | 0.75 | 0.02 |
| ENSRNOP00000055239 | AABR07060795.1 | 0.75 | 0.04 |
| ENSRNOP00000050806 | AABR07065823.2 | 0.75 | 0.00 |
| ENSRNOP00000016205 | Csad | 0.74 | 0.04 |
| ENSRNOP00000023485 | Ppic | 0.74 | 0.01 |
| ENSRNOP00000034048 | Ankrd46 | 0.74 | 0.04 |
| ENSRNOP00000031486 | AABR07034730.2 | 0.74 | 0.01 |
| ENSRNOP00000076111 | Nol8 | 0.74 | 0.03 |
| ENSRNOP00000018328 | Tgm2 | 0.74 | 0.05 |

| | | | |
|---|---|---|---|
| ENSRNOP00000070908 | Tmf1 | 0.74 | 0.01 |
| ENSRNOP00000073378 | Stim1 | 0.74 | 0.01 |
| ENSRNOP00000066948 | Pnkd | 0.73 | 0.00 |
| ENSRNOP00000075187 | Zfp326 | 0.73 | 0.03 |
| ENSRNOP00000004864 | Fmo3 | 0.73 | 0.01 |
| ENSRNOP00000073343 | Slc23a1 | 0.73 | 0.00 |
| ENSRNOP00000002699 | Sult1b1 | 0.73 | 0.03 |
| ENSRNOP00000003645 | Hgd | 0.73 | 0.00 |
| ENSRNOP00000066273 | AABR07065823.3 | 0.72 | 0.02 |
| ENSRNOP00000010288 | Fam151a | 0.71 | 0.01 |
| ENSRNOP00000032921 | Lemd3 | 0.71 | 0.01 |
| ENSRNOP00000039311 | LOC100365958 | 0.71 | 0.00 |
| ENSRNOP00000040223 | Hao2 | 0.71 | 0.04 |
| ENSRNOP00000074307 | Mme | 0.71 | 0.03 |
| ENSRNOP00000060620 | Tff3 | 0.70 | 0.00 |
| ENSRNOP00000024366 | Coa5 | 0.70 | 0.03 |
| ENSRNOP00000072936 | Ighm | 0.70 | 0.01 |
| ENSRNOP00000064877 | Hax1 | 0.69 | 0.00 |
| ENSRNOP00000072687 | AABR07061001.1 | 0.69 | 0.01 |
| ENSRNOP00000022074 | Fxyd2 | 0.69 | 0.01 |
| ENSRNOP00000024917 | Agt | 0.69 | 0.01 |
| ENSRNOP00000044411 | Gpt | 0.68 | 0.03 |
| ENSRNOP00000068562 | Slc2a5 | 0.68 | 0.00 |
| ENSRNOP00000029319 | AABR07060872.1 | 0.67 | 0.04 |
| ENSRNOP00000066134 | Zbtb33 | 0.67 | 0.02 |
| ENSRNOP00000065923 | Ggt1 | 0.67 | 0.01 |
| ENSRNOP00000013322 | Gale | 0.66 | 0.01 |
| ENSRNOP00000046491 | Hnrnpd | 0.66 | 0.01 |
| ENSRNOP00000035540 | Gclc | 0.66 | 0.01 |
| ENSRNOP00000017972 | Casp9 | 0.66 | 0.01 |
| ENSRNOP00000004956 | Col3a1 | 0.65 | 0.03 |
| ENSRNOP00000065896 | Dnah5 | 0.65 | 0.00 |
| ENSRNOP00000026462 | Psmb10 | 0.65 | 0.00 |
| ENSRNOP00000025529 | Atg4b | 0.65 | 0.03 |
| ENSRNOP00000007471 | Hnmt | 0.64 | 0.01 |
| ENSRNOP00000064447 | AC109901.2 | 0.64 | 0.03 |
| ENSRNOP00000022133 | Cep55 | 0.64 | 0.00 |
| ENSRNOP00000059125 | Akr1c12l1 | 0.63 | 0.00 |
| ENSRNOP00000064809 | Fabp5 | 0.62 | 0.00 |
| ENSRNOP00000023502 | Fam160a2 | 0.62 | 0.00 |
| ENSRNOP00000044037 | Igh-6 | 0.62 | 0.00 |
| ENSRNOP00000055191 | AABR07061005.1 | 0.62 | 0.03 |
| ENSRNOP00000003951 | Ren | 0.62 | 0.03 |
| ENSRNOP00000018343 | Gclm | 0.61 | 0.01 |
| ENSRNOP00000042340 | AABR07065886.1 | 0.60 | 0.01 |
| ENSRNOP00000067864 | Tbca | 0.60 | 0.01 |
| ENSRNOP00000072149 | AABR07065699.4 | 0.59 | 0.00 |
| ENSRNOP00000071041 | Cep89 | 0.59 | 0.01 |
| ENSRNOP00000070092 | AABR07065837.1 | 0.59 | 0.00 |
| ENSRNOP00000043834 | Smc1b | 0.58 | 0.01 |

| | | | |
|---|---|---|---|
| ENSRNOP00000065456 | Rad51ap2 | 0.58 | 0.00 |
| ENSRNOP00000032751 | Slc7a13 | 0.58 | 0.01 |
| ENSRNOP00000070616 | Atxn2l | 0.57 | 0.00 |
| ENSRNOP00000039261 | AABR07051684.1 | 0.56 | 0.01 |
| ENSRNOP00000055901 | Cryab | 0.55 | 0.01 |
| ENSRNOP00000071894 | Basp1 | 0.55 | 0.01 |
| ENSRNOP00000023037 | Itm2b | 0.54 | 0.02 |
| ENSRNOP00000022628 | Oat | 0.54 | 0.01 |
| ENSRNOP00000000026 | Slc21a4 | 0.54 | 0.04 |
| ENSRNOP00000010984 | Rgn | 0.53 | 0.01 |
| ENSRNOP00000040752 | Slco1a1 | 0.53 | 0.00 |
| ENSRNOP00000060692 | Anxa13 | 0.51 | 0.01 |
| ENSRNOP00000009581 | Slc3a1 | 0.49 | 0.00 |
| ENSRNOP00000020196 | Hp | 0.47 | 0.00 |
| ENSRNOP00000045662 | AABR07047899.1 | 0.45 | 0.05 |
| ENSRNOP00000067018 | LOC298111 | 0.37 | 0.02 |
| ENSRNOP00000068097 | LOC500473 | 0.30 | 0.02 |

**Supplemental Table S3.** Differentially expressed proteins in the Sch-treated vs. control group.

| Accession | Gene Name | S/C | t test p value |
|---|---|---|---|
| ENSRNOP00000041060 | Mt1m | 3.93 | 0.01 |
| ENSRNOP00000026122 | Hmgcs2 | 3.60 | 0.04 |
| ENSRNOP00000054706 | Mt1 | 2.67 | 0.02 |
| ENSRNOP00000028664 | Mllt11 | 2.23 | 0.00 |
| ENSRNOP00000066577 | Gpx1 | 1.97 | 0.01 |
| ENSRNOP00000023430 | Cep192 | 1.89 | 0.02 |
| ENSRNOP00000060140 | Acaa2 | 1.75 | 0.04 |
| ENSRNOP00000011784 | Eci1 | 1.72 | 0.03 |
| ENSRNOP00000065846 | Wwc3 | 1.71 | 0.02 |
| ENSRNOP00000058419 | Bhmt2 | 1.70 | 0.05 |
| ENSRNOP00000002410 | Ehhadh | 1.68 | 0.01 |
| ENSRNOP00000056322 | Ankrd33b | 1.63 | 0.00 |
| ENSRNOP00000054540 | Ttc39a | 1.62 | 0.00 |
| ENSRNOP00000018622 | Rbp1 | 1.61 | 0.02 |
| ENSRNOP00000073777 | Myo1g | 1.60 | 0.00 |
| ENSRNOP00000055007 | Tsc22d1 | 1.60 | 0.02 |
| ENSRNOP00000024622 | LOC501233 | 1.58 | 0.00 |
| ENSRNOP00000011945 | Gpnmb | 1.58 | 0.01 |
| ENSRNOP00000037803 | Fth1 | 1.58 | 0.01 |
| ENSRNOP00000041451 | Hba-a3 | 1.57 | 0.02 |
| ENSRNOP00000017712 | Nudt2 | 1.57 | 0.01 |
| ENSRNOP00000008613 | Slc5a8 | 1.57 | 0.01 |
| ENSRNOP00000065206 | Apoa1 | 1.57 | 0.05 |
| ENSRNOP00000005622 | S100g | 1.56 | 0.01 |
| ENSRNOP00000075331 | Ddr1 | 1.55 | 0.00 |
| ENSRNOP00000023739 | Akr1c3 | 1.54 | 0.00 |
| ENSRNOP00000068322 | Tmlhe | 1.53 | 0.01 |
| ENSRNOP00000022150 | Cd1d1 | 1.53 | 0.01 |
| ENSRNOP00000002712 | Ugt2b35 | 1.53 | 0.01 |
| ENSRNOP00000054044 | Naglt1 | 1.52 | 0.00 |
| ENSRNOP00000018341 | Myd88 | 1.51 | 0.02 |
| ENSRNOP00000030913 | Pck1 | 1.51 | 0.00 |
| ENSRNOP00000010760 | Mettl7b | 1.48 | 0.02 |
| ENSRNOP00000066155 | Amt | 1.48 | 0.00 |
| ENSRNOP00000051846 | Slc25a10 | 1.48 | 0.03 |
| ENSRNOP00000055038 | Gstt3 | 1.47 | 0.03 |
| ENSRNOP00000027537 | Ech1 | 1.47 | 0.01 |
| ENSRNOP00000075208 | Kb23 | 1.46 | 0.03 |
| ENSRNOP00000014213 | Acot7 | 1.45 | 0.01 |
| ENSRNOP00000075693 | LOC299282 | 1.44 | 0.03 |
| ENSRNOP00000016784 | Trio | 1.44 | 0.00 |
| ENSRNOP00000002528 | Far2 | 1.44 | 0.01 |
| ENSRNOP00000008337 | Aco1 | 1.43 | 0.00 |
| ENSRNOP00000021694 | Vnn1 | 1.42 | 0.02 |
| ENSRNOP00000073662 | Pigr | 1.42 | 0.03 |
| ENSRNOP00000008680 | Tmem70 | 1.42 | 0.02 |
| ENSRNOP00000024000 | Aldh1a1 | 1.42 | 0.00 |

| | | | |
|---|---|---|---|
| ENSRNOP00000073009 | Krt34 | 1.41 | 0.02 |
| ENSRNOP00000043298 | Cyp24a1 | 1.41 | 0.00 |
| ENSRNOP00000005641 | Pdk2 | 1.41 | 0.00 |
| ENSRNOP00000012759 | Pdk4 | 1.40 | 0.01 |
| ENSRNOP00000001975 | Baz1b | 1.40 | 0.00 |
| ENSRNOP00000068006 | Stmn1 | 1.39 | 0.01 |
| ENSRNOP00000005551 | Adipor1 | 1.39 | 0.00 |
| ENSRNOP00000038073 | Hadha | 1.38 | 0.00 |
| ENSRNOP00000028098 | Rnls | 1.37 | 0.02 |
| ENSRNOP00000074978 | Hmgn2 | 1.37 | 0.04 |
| ENSRNOP00000044955 | Paqr9 | 1.37 | 0.03 |
| ENSRNOP00000072623 | LOC681544 | 1.37 | 0.01 |
| ENSRNOP00000060618 | Rabl6 | 1.37 | 0.00 |
| ENSRNOP00000040341 | Apon | 1.37 | 0.02 |
| ENSRNOP00000016445 | Hddc3 | 1.37 | 0.01 |
| ENSRNOP00000061162 | RGD1565355 | 1.37 | 0.02 |
| ENSRNOP00000022713 | Itga8 | 1.36 | 0.01 |
| ENSRNOP00000015576 | Akap2 | 1.36 | 0.00 |
| ENSRNOP00000025859 | Myh7b | 1.36 | 0.00 |
| ENSRNOP00000065593 | Ces2c | 1.36 | 0.01 |
| ENSRNOP00000069993 | Ptms | 1.36 | 0.03 |
| ENSRNOP00000043148 | Comt | 1.36 | 0.00 |
| ENSRNOP00000065713 | Akr1b8 | 1.36 | 0.00 |
| ENSRNOP00000004866 | Jchain | 1.36 | 0.03 |
| ENSRNOP00000001365 | Ccz1b | 1.36 | 0.01 |
| ENSRNOP00000016954 | Cpt2 | 1.33 | 0.01 |
| ENSRNOP00000014637 | Hadhb | 1.33 | 0.00 |
| ENSRNOP00000051538 | Acox1 | 1.33 | 0.02 |
| ENSRNOP00000071112 | Gstt2 | 1.32 | 0.01 |
| ENSRNOP00000024196 | Gpld1 | 1.32 | 0.00 |
| ENSRNOP00000012036 | Gnpnat1 | 1.32 | 0.03 |
| ENSRNOP00000060590 | Naga | 1.31 | 0.00 |
| ENSRNOP00000017468 | Ldha | 1.31 | 0.02 |
| ENSRNOP00000013754 | Acot4 | 1.30 | 0.03 |
| ENSRNOP00000002962 | Mrps18c | 1.30 | 0.04 |
| ENSRNOP00000026812 | Actn3 | 1.30 | 0.00 |
| ENSRNOP00000065600 | Ahsa1 | 1.29 | 0.00 |
| ENSRNOP00000040878 | LOC108351137 | 1.29 | 0.02 |
| ENSRNOP00000071799 | Slc43a3 | 1.29 | 0.03 |
| ENSRNOP00000057148 | Adtrp | 1.29 | 0.00 |
| ENSRNOP00000023984 | Itih3 | 1.29 | 0.01 |
| ENSRNOP00000016372 | Glrx | 1.29 | 0.00 |
| ENSRNOP00000014267 | Car1 | 1.28 | 0.02 |
| ENSRNOP00000041926 | Slc5a12 | 1.28 | 0.01 |
| ENSRNOP00000073940 | Pcdh1 | 1.28 | 0.04 |
| ENSRNOP00000003274 | Serpinb8 | 1.28 | 0.00 |
| ENSRNOP00000031836 | Yod1 | 1.27 | 0.02 |
| ENSRNOP00000042132 | Acox1 | 1.27 | 0.01 |
| ENSRNOP00000039445 | Espn | 1.27 | 0.03 |
| ENSRNOP00000005204 | Stx8 | 1.27 | 0.02 |

| | | | |
|---|---|---|---|
| ENSRNOP00000001960 | Tmem120a | 1.26 | 0.01 |
| ENSRNOP00000010602 | Aup1 | 1.26 | 0.02 |
| ENSRNOP00000008504 | Hspa2 | 1.26 | 0.03 |
| ENSRNOP00000066485 | Bola2 | 1.26 | 0.01 |
| ENSRNOP00000072136 | Gpx2 | 1.26 | 0.03 |
| ENSRNOP00000022022 | Eci3 | 1.26 | 0.02 |
| ENSRNOP00000017686 | Acadl | 1.26 | 0.03 |
| ENSRNOP00000013262 | Etfdh | 1.26 | 0.04 |
| ENSRNOP00000069919 | Tagln | 1.26 | 0.04 |
| ENSRNOP00000026564 | Mthfd1l | 1.25 | 0.05 |
| ENSRNOP00000049160 | Stradb | 1.25 | 0.03 |
| ENSRNOP00000000494 | Fkbpl | 1.25 | 0.03 |
| ENSRNOP00000001277 | Slc15a4 | 1.25 | 0.00 |
| ENSRNOP00000027520 | Slc25a20 | 1.25 | 0.05 |
| ENSRNOP00000056213 | Hsdl2 | 1.25 | 0.03 |
| ENSRNOP00000007365 | Coch | 1.24 | 0.00 |
| ENSRNOP00000026651 | Ubxn1 | 1.24 | 0.00 |
| ENSRNOP00000072116 | Clu | 1.24 | 0.01 |
| ENSRNOP00000067980 | RGD1565355 | 1.24 | 0.02 |
| ENSRNOP00000036574 | Acad11 | 1.24 | 0.01 |
| ENSRNOP00000013703 | Enpp5 | 1.24 | 0.03 |
| ENSRNOP00000014268 | Nrg1 | 1.24 | 0.01 |
| ENSRNOP00000009960 | Sorcs2 | 1.24 | 0.01 |
| ENSRNOP00000053510 | Gp1ba | 1.24 | 0.02 |
| ENSRNOP00000003004 | Hsd17b11 | 1.24 | 0.00 |
| ENSRNOP00000025534 | C5 | 1.23 | 0.01 |
| ENSRNOP00000069382 | Slco2a1 | 1.23 | 0.00 |
| ENSRNOP00000009627 | Acss1 | 1.23 | 0.03 |
| ENSRNOP00000024659 | Slc27a1 | 1.22 | 0.00 |
| ENSRNOP00000028194 | Dhcr7 | 1.22 | 0.01 |
| ENSRNOP00000018545 | C9 | 1.22 | 0.04 |
| ENSRNOP00000038628 | Dgka | 1.22 | 0.04 |
| ENSRNOP00000039630 | Bin2 | 1.22 | 0.01 |
| ENSRNOP00000005459 | Marc1 | 1.22 | 0.04 |
| ENSRNOP00000012888 | Cyp4a1 | 1.22 | 0.02 |
| ENSRNOP00000070446 | Vdr | 1.22 | 0.02 |
| ENSRNOP00000072412 | Mrpl14 | 1.22 | 0.04 |
| ENSRNOP00000030271 | Mgll | 1.22 | 0.00 |
| ENSRNOP00000048774 | RT1-Da | 1.22 | 0.04 |
| ENSRNOP00000007927 | Rpia | 1.22 | 0.03 |
| ENSRNOP00000027925 | Slc22a18 | 1.22 | 0.04 |
| ENSRNOP00000006521 | Mta1 | 1.22 | 0.03 |
| ENSRNOP00000029498 | Fastkd3 | 1.21 | 0.05 |
| ENSRNOP00000002864 | Gtf3c3 | 1.21 | 0.02 |
| ENSRNOP00000016948 | Bag2 | 1.21 | 0.00 |
| ENSRNOP00000071497 | Tinag | 1.21 | 0.01 |
| ENSRNOP00000001077 | RGD1302996 | 1.21 | 0.02 |
| ENSRNOP00000037398 | Yars2 | 1.21 | 0.01 |
| ENSRNOP00000038337 | Iba57 | 1.21 | 0.01 |
| ENSRNOP00000021545 | Plgrkt | 1.21 | 0.04 |

| | | | |
|---|---|---|---|
| ENSRNOP00000074700 | Uqcr10 | 1.21 | 0.01 |
| ENSRNOP00000074322 | Kif1b | 1.21 | 0.00 |
| ENSRNOP00000012757 | Papln | 1.21 | 0.03 |
| ENSRNOP00000064884 | Ces2c | 1.21 | 0.04 |
| ENSRNOP00000024648 | Arl6ip1 | 1.20 | 0.03 |
| ENSRNOP00000033445 | Tcerg1l | 1.20 | 0.04 |
| ENSRNOP00000000170 | Atg12 | 1.20 | 0.03 |
| ENSRNOP00000064332 | Tsfm | 1.20 | 0.03 |
| ENSRNOP00000010763 | Pxk | 0.83 | 0.02 |
| ENSRNOP00000050393 | Nf1 | 0.83 | 0.02 |
| ENSRNOP00000043362 | Mup4 | 0.83 | 0.01 |
| ENSRNOP00000034657 | LOC100360647 | 0.83 | 0.02 |
| ENSRNOP00000022986 | Mindy3 | 0.83 | 0.01 |
| ENSRNOP00000065771 | Lrg1 | 0.83 | 0.03 |
| ENSRNOP00000070394 | Fdps | 0.83 | 0.05 |
| ENSRNOP00000003282 | Gpc4 | 0.83 | 0.01 |
| ENSRNOP00000075216 | Clec16a | 0.83 | 0.00 |
| ENSRNOP00000071689 | Cand2 | 0.83 | 0.01 |
| ENSRNOP00000066553 | LOC684270 | 0.83 | 0.00 |
| ENSRNOP00000037089 | Slco4c1 | 0.83 | 0.02 |
| ENSRNOP00000027999 | Acadsb | 0.83 | 0.01 |
| ENSRNOP00000064116 | Klhdc7a | 0.83 | 0.02 |
| ENSRNOP00000072188 | Slc7a8 | 0.83 | 0.03 |
| ENSRNOP00000024464 | Clic4 | 0.83 | 0.04 |
| ENSRNOP00000063118 | Gipc2 | 0.83 | 0.05 |
| ENSRNOP00000007374 | Hpcal1 | 0.83 | 0.00 |
| ENSRNOP00000071707 | Slc39a7 | 0.83 | 0.01 |
| ENSRNOP00000001228 | Psph | 0.83 | 0.00 |
| ENSRNOP00000033324 | Golga7 | 0.83 | 0.01 |
| ENSRNOP00000074171 | Cacna2d1 | 0.83 | 0.05 |
| ENSRNOP00000003092 | Casr | 0.82 | 0.01 |
| ENSRNOP00000004525 | Cenpf | 0.82 | 0.00 |
| ENSRNOP00000027212 | Nfkbib | 0.82 | 0.03 |
| ENSRNOP00000045388 | AABR07044383.1 | 0.82 | 0.05 |
| ENSRNOP00000032681 | Tmem126a | 0.82 | 0.00 |
| ENSRNOP00000017139 | Pcbp4 | 0.82 | 0.01 |
| ENSRNOP00000003189 | Slc15a2 | 0.82 | 0.01 |
| ENSRNOP00000044552 | Rela | 0.82 | 0.00 |
| ENSRNOP00000006597 | Slc25a35 | 0.82 | 0.01 |
| ENSRNOP00000014496 | RGD1309534 | 0.82 | 0.00 |
| ENSRNOP00000071311 | Folr1 | 0.82 | 0.01 |
| ENSRNOP00000039968 | Cbs | 0.81 | 0.00 |
| ENSRNOP00000049903 | Ggct | 0.81 | 0.02 |
| ENSRNOP00000016504 | Abhd14b | 0.81 | 0.02 |
| ENSRNOP00000043021 | AABR07044570.1 | 0.81 | 0.02 |
| ENSRNOP00000048749 | Rap1gap | 0.81 | 0.01 |
| ENSRNOP00000018646 | Snrpd1 | 0.81 | 0.03 |
| ENSRNOP00000013845 | Dennd10 | 0.81 | 0.00 |
| ENSRNOP00000003237 | Prkg2 | 0.81 | 0.01 |
| ENSRNOP00000005100 | Slc43a2 | 0.81 | 0.02 |

| | | | |
|---|---|---|---|
| ENSRNOP00000064959 | AABR07050399.1 | 0.81 | 0.02 |
| ENSRNOP00000073635 | Smurf1 | 0.81 | 0.04 |
| ENSRNOP00000061340 | Slc2a1 | 0.81 | 0.00 |
| ENSRNOP00000043492 | AC107446.2 | 0.81 | 0.01 |
| ENSRNOP00000075613 | Arhgap24 | 0.81 | 0.00 |
| ENSRNOP00000021797 | Hars2 | 0.81 | 0.03 |
| ENSRNOP00000018170 | Mat2a | 0.81 | 0.00 |
| ENSRNOP00000068773 | Dnajb14 | 0.80 | 0.00 |
| ENSRNOP00000025196 | Slc3a2 | 0.80 | 0.02 |
| ENSRNOP00000067214 | Zfpl1 | 0.80 | 0.01 |
| ENSRNOP00000023274 | Cdk11b | 0.80 | 0.00 |
| ENSRNOP00000004864 | Fmo3 | 0.80 | 0.00 |
| ENSRNOP00000002540 | Sdf2l1 | 0.80 | 0.02 |
| ENSRNOP00000023383 | Vac14 | 0.80 | 0.03 |
| ENSRNOP00000026597 | Pgpep1 | 0.80 | 0.02 |
| ENSRNOP00000071202 | AABR07060886.1 | 0.80 | 0.02 |
| ENSRNOP00000073742 | Rufy3 | 0.80 | 0.04 |
| ENSRNOP00000015888 | Vill | 0.80 | 0.02 |
| ENSRNOP00000028912 | Dusp23 | 0.80 | 0.02 |
| ENSRNOP00000048676 | Atp11c | 0.80 | 0.00 |
| ENSRNOP00000073343 | Slc23a1 | 0.80 | 0.02 |
| ENSRNOP00000027220 | Scly | 0.79 | 0.00 |
| ENSRNOP00000075159 | Lrp2 | 0.79 | 0.00 |
| ENSRNOP00000068163 | Ly6e | 0.79 | 0.05 |
| ENSRNOP00000003713 | Npl | 0.79 | 0.02 |
| ENSRNOP00000038185 | Slc13a2 | 0.79 | 0.04 |
| ENSRNOP00000068285 | AABR07048475.1 | 0.79 | 0.02 |
| ENSRNOP00000068834 | Picalm | 0.79 | 0.01 |
| ENSRNOP00000006917 | Atp6v1c1 | 0.79 | 0.00 |
| ENSRNOP00000005844 | Pah | 0.79 | 0.00 |
| ENSRNOP00000044335 | Nr3c1 | 0.79 | 0.01 |
| ENSRNOP00000073179 | Ilf3 | 0.79 | 0.01 |
| ENSRNOP00000023485 | Ppic | 0.78 | 0.01 |
| ENSRNOP00000062936 | AABR07065792.1 | 0.78 | 0.02 |
| ENSRNOP00000027737 | Mrpl54 | 0.78 | 0.05 |
| ENSRNOP00000011237 | Plpp3 | 0.78 | 0.01 |
| ENSRNOP00000070467 | Slc5a9 | 0.78 | 0.00 |
| ENSRNOP00000001941 | Polr2j | 0.78 | 0.00 |
| ENSRNOP00000039536 | Ighm | 0.78 | 0.02 |
| ENSRNOP00000072092 | Dpp4 | 0.78 | 0.03 |
| ENSRNOP00000020192 | Hagh | 0.78 | 0.00 |
| ENSRNOP00000064877 | Hax1 | 0.78 | 0.02 |
| ENSRNOP00000018837 | Wwp2 | 0.77 | 0.01 |
| ENSRNOP00000015692 | Aqp1 | 0.77 | 0.04 |
| ENSRNOP00000051747 | Lsp1 | 0.77 | 0.01 |
| ENSRNOP00000000481 | Skiv2l | 0.77 | 0.01 |
| ENSRNOP00000028887 | Pcna | 0.77 | 0.00 |
| ENSRNOP00000031564 | Tubal3 | 0.77 | 0.01 |
| ENSRNOP00000023080 | Qprt | 0.77 | 0.02 |
| ENSRNOP00000073116 | Asah2 | 0.77 | 0.00 |

| | | | |
|---|---|---|---|
| ENSRNOP00000070222 | Ttc39c | 0.77 | 0.04 |
| ENSRNOP00000042580 | AABR07065768.1 | 0.77 | 0.01 |
| ENSRNOP00000031764 | Slc34a1 | 0.77 | 0.00 |
| ENSRNOP00000029808 | Fdft1 | 0.77 | 0.00 |
| ENSRNOP00000028828 | Mrps26 | 0.76 | 0.03 |
| ENSRNOP00000073378 | Stim1 | 0.76 | 0.01 |
| ENSRNOP00000055169 | Akr1c1 | 0.76 | 0.00 |
| ENSRNOP00000066948 | Pnkd | 0.76 | 0.01 |
| ENSRNOP00000063373 | Ccdc88a | 0.76 | 0.03 |
| ENSRNOP00000003645 | Hgd | 0.76 | 0.00 |
| ENSRNOP00000004298 | Anks3 | 0.76 | 0.00 |
| ENSRNOP00000019620 | Rer1 | 0.76 | 0.04 |
| ENSRNOP00000071870 | Sat2 | 0.75 | 0.00 |
| ENSRNOP00000011384 | Mthfr | 0.75 | 0.01 |
| ENSRNOP00000068562 | Slc2a5 | 0.75 | 0.05 |
| ENSRNOP00000006131 | Atg2b | 0.75 | 0.04 |
| ENSRNOP00000004867 | Sumo2 | 0.75 | 0.02 |
| ENSRNOP00000018003 | Hykk | 0.75 | 0.00 |
| ENSRNOP00000026726 | Wdr3 | 0.75 | 0.04 |
| ENSRNOP00000048768 | Myl10 | 0.74 | 0.02 |
| ENSRNOP00000072904 | Arhgef1 | 0.74 | 0.04 |
| ENSRNOP00000021657 | Pccb | 0.74 | 0.01 |
| ENSRNOP00000016912 | Mgam | 0.74 | 0.00 |
| ENSRNOP00000034048 | Ankrd46 | 0.74 | 0.04 |
| ENSRNOP00000005440 | Tcaim | 0.74 | 0.02 |
| ENSRNOP00000005262 | Cpd | 0.73 | 0.04 |
| ENSRNOP00000019571 | Retsat | 0.73 | 0.05 |
| ENSRNOP00000076111 | Nol8 | 0.73 | 0.01 |
| ENSRNOP00000001508 | Prkab1 | 0.73 | 0.02 |
| ENSRNOP00000045992 | Myl12b | 0.73 | 0.01 |
| ENSRNOP00000075187 | Zfp326 | 0.72 | 0.03 |
| ENSRNOP00000076301 | Ass1 | 0.72 | 0.00 |
| ENSRNOP00000031486 | AABR07034730.2 | 0.71 | 0.01 |
| ENSRNOP00000042202 | Siae | 0.71 | 0.03 |
| ENSRNOP00000023211 | Clptm1l | 0.71 | 0.04 |
| ENSRNOP00000066421 | AABR07065776.3 | 0.70 | 0.00 |
| ENSRNOP00000032921 | Lemd3 | 0.70 | 0.01 |
| ENSRNOP00000072687 | AABR07061001.1 | 0.70 | 0.02 |
| ENSRNOP00000024366 | Coa5 | 0.70 | 0.03 |
| ENSRNOP00000066331 | AABR07065750.2 | 0.70 | 0.05 |
| ENSRNOP00000022113 | Ttr | 0.70 | 0.00 |
| ENSRNOP00000010288 | Fam151a | 0.69 | 0.01 |
| ENSRNOP00000060620 | Tff3 | 0.69 | 0.00 |
| ENSRNOP00000050806 | AABR07065823.2 | 0.69 | 0.00 |
| ENSRNOP00000072936 | Ighm | 0.68 | 0.00 |
| ENSRNOP00000010717 | Frmd4b | 0.68 | 0.01 |
| ENSRNOP00000064809 | Fabp5 | 0.68 | 0.01 |
| ENSRNOP00000024917 | Agt | 0.68 | 0.01 |
| ENSRNOP00000066273 | AABR07065823.3 | 0.68 | 0.01 |
| ENSRNOP00000070120 | AABR07060963.2 | 0.68 | 0.00 |

| | | | |
|---|---|---|---|
| ENSRNOP00000072454 | AABR07065714.1 | 0.67 | 0.00 |
| ENSRNOP00000059125 | Akr1c12l1 | 0.67 | 0.00 |
| ENSRNOP00000035540 | Gclc | 0.67 | 0.00 |
| ENSRNOP00000065923 | Ggt1 | 0.67 | 0.01 |
| ENSRNOP00000046491 | Hnrnpd | 0.66 | 0.03 |
| ENSRNOP00000064447 | AC109901.2 | 0.66 | 0.03 |
| ENSRNOP00000067864 | Tbca | 0.66 | 0.02 |
| ENSRNOP00000007471 | Hnmt | 0.65 | 0.00 |
| ENSRNOP00000072149 | AABR07065699.4 | 0.65 | 0.00 |
| ENSRNOP00000026462 | Psmb10 | 0.65 | 0.00 |
| ENSRNOP00000025529 | Atg4b | 0.65 | 0.04 |
| ENSRNOP00000074126 | LOC100365958 | 0.65 | 0.01 |
| ENSRNOP00000070616 | Atxn2l | 0.64 | 0.00 |
| ENSRNOP00000049378 | Pcid2 | 0.64 | 0.00 |
| ENSRNOP00000065896 | Dnah5 | 0.64 | 0.01 |
| ENSRNOP00000042340 | AABR07065886.1 | 0.63 | 0.00 |
| ENSRNOP00000013322 | Gale | 0.63 | 0.02 |
| ENSRNOP00000044037 | Igh-6 | 0.63 | 0.00 |
| ENSRNOP00000029319 | AABR07060872.1 | 0.63 | 0.02 |
| ENSRNOP00000018343 | Gclm | 0.62 | 0.00 |
| ENSRNOP00000003951 | Ren | 0.61 | 0.01 |
| ENSRNOP00000043834 | Smc1b | 0.61 | 0.03 |
| ENSRNOP00000023502 | Fam160a2 | 0.61 | 0.00 |
| ENSRNOP00000039311 | LOC100365958 | 0.61 | 0.00 |
| ENSRNOP00000065456 | Rad51ap2 | 0.60 | 0.00 |
| ENSRNOP00000032751 | Slc7a13 | 0.60 | 0.01 |
| ENSRNOP00000022133 | Cep55 | 0.58 | 0.00 |
| ENSRNOP00000022628 | Oat | 0.58 | 0.02 |
| ENSRNOP00000017972 | Casp9 | 0.58 | 0.00 |
| ENSRNOP00000066134 | Zbtb33 | 0.58 | 0.01 |
| ENSRNOP00000040752 | Slco1a1 | 0.57 | 0.00 |
| ENSRNOP00000010984 | Rgn | 0.55 | 0.00 |
| ENSRNOP00000071041 | Cep89 | 0.55 | 0.01 |
| ENSRNOP00000070092 | AABR07065837.1 | 0.54 | 0.00 |
| ENSRNOP00000071894 | Basp1 | 0.53 | 0.01 |
| ENSRNOP00000039261 | AABR07051684.1 | 0.53 | 0.01 |
| ENSRNOP00000055191 | AABR07061005.1 | 0.52 | 0.01 |
| ENSRNOP00000060692 | Anxa13 | 0.52 | 0.01 |
| ENSRNOP00000009581 | Slc3a1 | 0.48 | 0.00 |
| ENSRNOP00000023037 | Itm2b | 0.41 | 0.00 |
| ENSRNOP00000067018 | LOC298111 | 0.40 | 0.04 |
| ENSRNOP00000023385 | Ephx2 | 0.38 | 0.01 |
| ENSRNOP00000068097 | LOC500473 | 0.30 | 0.03 |

**Supplemental Table S4.** Differentially expressed proteins in the Sch-treated vs. model group.

| Accession | Gene Name | S/M | t test p value |
|---|---|---|---|
| ENSRNOP00000058825 | Pvalb | 1.62 | 0.01 |
| ENSRNOP00000022150 | Cd1d1 | 1.48 | 0.01 |
| ENSRNOP00000016130 | Psip1 | 1.42 | 0.05 |
| ENSRNOP00000056322 | Ankrd33b | 1.41 | 0.02 |
| ENSRNOP00000018341 | Myd88 | 1.39 | 0.04 |
| ENSRNOP00000073009 | Krt34 | 1.33 | 0.03 |
| ENSRNOP00000065414 | AABR07030903.1 | 1.29 | 0.00 |
| ENSRNOP00000071799 | Slc43a3 | 1.27 | 0.04 |
| ENSRNOP00000027986 | Eif3g | 1.27 | 0.02 |
| ENSRNOP00000072449 | Bgn | 1.26 | 0.03 |
| ENSRNOP00000008504 | Hspa2 | 1.25 | 0.04 |
| ENSRNOP00000053391 | Xaf1 | 1.24 | 0.05 |
| ENSRNOP00000026812 | Actn3 | 1.23 | 0.01 |
| ENSRNOP00000003237 | Prkg2 | 0.83 | 0.02 |
| ENSRNOP00000000481 | Skiv2l | 0.82 | 0.00 |
| ENSRNOP00000071249 | Zc3h12c | 0.82 | 0.01 |
| ENSRNOP00000026225 | Ifi30 | 0.82 | 0.00 |
| ENSRNOP00000003713 | Npl | 0.82 | 0.03 |
| ENSRNOP00000037197 | Slc35d1 | 0.82 | 0.03 |
| ENSRNOP00000001941 | Polr2j | 0.82 | 0.00 |
| ENSRNOP00000005129 | Tmigd1 | 0.82 | 0.04 |
| ENSRNOP00000073179 | Ilf3 | 0.82 | 0.01 |
| ENSRNOP00000004867 | Sumo2 | 0.81 | 0.03 |
| ENSRNOP00000070149 | Srpk2 | 0.80 | 0.03 |
| ENSRNOP00000028887 | Pcna | 0.80 | 0.00 |
| ENSRNOP00000049378 | Pcid2 | 0.80 | 0.00 |
| ENSRNOP00000064452 | Crb3 | 0.80 | 0.01 |
| ENSRNOP00000020401 | Tent5c | 0.79 | 0.03 |
| ENSRNOP00000064959 | AABR07050399.1 | 0.79 | 0.02 |
| ENSRNOP00000035928 | Nmi | 0.79 | 0.04 |
| ENSRNOP00000019620 | Rer1 | 0.79 | 0.04 |
| ENSRNOP00000005100 | Slc43a2 | 0.79 | 0.03 |
| ENSRNOP00000018646 | Snrpd1 | 0.78 | 0.01 |
| ENSRNOP00000011237 | Plpp3 | 0.78 | 0.01 |
| ENSRNOP00000021797 | Hars2 | 0.77 | 0.02 |
| ENSRNOP00000066775 | Tspan4 | 0.77 | 0.01 |
| ENSRNOP00000070222 | Ttc39c | 0.76 | 0.03 |
| ENSRNOP00000061900 | Bag4 | 0.75 | 0.02 |
| ENSRNOP00000049566 | Nckipsd | 0.74 | 0.02 |
| ENSRNOP00000026726 | Wdr3 | 0.74 | 0.04 |
| ENSRNOP00000006658 | Zfp512 | 0.73 | 0.02 |
| ENSRNOP00000010717 | Frmd4b | 0.72 | 0.01 |
| ENSRNOP00000013369 | Wdr45 | 0.72 | 0.01 |
| ENSRNOP00000031564 | Tubal3 | 0.71 | 0.01 |
| ENSRNOP00000001508 | Prkab1 | 0.66 | 0.01 |

| | | | |
|---|---|---|---|
| ENSRNOP00000009136 | Vcpip1 | 0.62 | 0.03 |
