## Supplementary material for "Quantitative proteomics combined with network pharmacology analysis unveils the biological basis of Schisandrin B in treating diabetic nephropathy": Affidavit of Approval of Animal Ethical and Welfare

### 天津大学动物实验伦理审查证明

#### Affidavit of Approval of Animal Ethical and Welfare of Tianjin University

| 编号 Approval No. | TJUE-2022-009 |
|---|---|

以下《动物实验方案》经过实验动物伦理委员会审核，符合动物保护、动物福利和伦理原则，符合国家实验动物福利伦理的相关规定，特此证明。

The animal use protocol listed below has been reviewed and approved by the Animal Ethical and Welfare Committee (AEWC), Hereby certify.

**审查类型：** ☐ 项目申报 ■ 动物实验

**递交材料：** ■ 动物福利伦理审查申请表

<table>
<tr><td rowspan="2">项 目 名 称<br>Protocol Title</td><td colspan="4">定量蛋白质组学结合网络药理学分析揭示五味子乙素治疗糖尿病肾病的生物学基础</td></tr>
<tr><td colspan="4">Quantitative proteomics combined with network pharmacology analysis to unveil the biological basis of Schisandrin B in treating Diabetic Nephropathy</td></tr>
<tr><td>Kang Jun</td><td colspan="2">;<br></td></tr>
<tr><td>Kang Jun</td><td colspan="2">;<br></td></tr>
<tr><td rowspan="2">院系<br>Department</td><td colspan="2">生命科学学院</td><td rowspan="2">申 请 日 期<br>Application date</td><td rowspan="2">Jan 21, 2022</td></tr>
<tr><td colspan="2">School of life sciences</td></tr>
<tr><td rowspan="2">动物种系<br>Species or Strains</td><td colspan="2">SD 大鼠</td><td rowspan="2">动 物 数 量<br>Quantity</td><td rowspan="2">50</td></tr>
<tr><td colspan="2">SD rats</td></tr>
<tr><td colspan="5">实验动物使用许可证号 Number of Animal use permit:<br>No. 110322210100331916</td></tr>
<tr><td colspan="5">审查意见 Results of inspection:<br>该项目符合动物福利伦理相关要求（Agree）。<br>负责人 Chief Officer: [signature]<br>日期 Date: 2022/2/11<br>签章（实验动物伦理委员会）Stamp</td></tr>
</table>
