## Supplementary material for "Quantitative proteomics combined with network pharmacology analysis unveils the biological basis of Schisandrin B in treating diabetic nephropathy": Certificate of English Language Editing

### Manuscript Title:

### Date of Revision:

September 20, 2022


Abstract:

Background: Diabetic nephropathy (DN) is a major complication of diabetes, leading to end-stage renal failure in severe cases. Schisandrin B (Sch) is a natural pharmaceutical monomer that was shown to prevent kidney damage caused by diabetes and to restore its function. However, there is still a lack of comprehensive and systematic understanding of the mechanism of Sch treatment in DN.

Purpose: We aim to provide a systematic overview of the mechanisms of Sch in multiple pathways to treat DN in rats.

Methods: Streptozocin was used to build a DN rat model. The DN rats were further treated with Sch. The possible mechanism of Sch protective effects against DN was predicted using network pharmacology and was verified by a quantitative proteomics analysis.

Results: High dose Sch treatment significantly downregulated fasting blood glucose, creatinine, blood urea nitrogen, and urinary protein levels and reduced collagen deposition in the glomeruli...


This document certifies that the manuscript listed above was copy edited for English language by LetPub, with regard to grammar, punctuation, spelling, and clarity. All of our language editors are native English speakers with long-term experience in editing scientific and technical manuscripts. We are committed to leveling the playing field for researchers whose native language is not English.

- Documents receiving this certification should be regarded as having undergone professional editorial revision for English language before submission. However, the authors may accept or reject LetPub's suggestions and changes at their own discretion and LetPub does not have editorial control over the submitted documents.
- The language quality of the submitted document is the sole responsibility of the submitting authors subject to those authors' adherence to LetPub's revisions and instruction. LetPub's provision of service does not constitute a guarantee or endorsement of the authors' work herein.
- Neither the research content nor the authors' intended meaning were altered in any way during the editing process.
- If you have any questions or concerns about this edited document, please contact us at

LetPub
Your Manuscript Editor

LetPub is an author service brand owned and operated by Accdon LLC. Headquartered in the Boston area, we are a full-spectrum author services company with a large team of US-based certified language and scientific editors, ISO 17001 accredited translators, and professional scientific illustrators and animators. We advocate ethical publication practices and are an official member of the Committee on Publication Ethics (COPE).

For more information about our company, services, and partnership programs, please visit www.letpub.com.


© 2022 Accdon, LLC. All Rights Reserved.

Tel: 1-781-202-9968 Address: 400 Fifth Ave, Suite 530, Waltham, MA 02451, United States
